# Analysis of conditional colocalization relationships and hierarchies in three-color microscopy images

**DOI:** 10.1101/2021.06.16.448703

**Authors:** Jesus Vega-Lugo, Bruno da Rocha-Azevedo, Aparajita Dasgupta, Nicolas Touret, Khuloud Jaqaman

## Abstract

Colocalization analysis of multicolor microscopy images is a cornerstone approach in cell biology. It provides information on the localization of molecules within various subcellular compartments and allows the interrogation of molecular interactions in their cellular context. However, almost all colocalization analyses are designed for two-color images. This limits their applicability and the type of information that they reveal, leading to underutilization of multicolor microscopy images. Here we describe an approach, termed “conditional colocalization analysis,” for analyzing the colocalization relationships between three molecular entities in three-color microscopy images. Going beyond the question of whether colocalization is present or not, it addresses the question of whether the colocalization between two entities is influenced, positively or negatively, by their colocalization with a third entity. We showcase two applications of conditional colocalization analysis, one addressing the question of compartmentalization of molecular interactions, and one investigating the hierarchy of molecular interactions in a multimolecular complex. The software for conditional colocalization analysis is available at https://github.com/kjaqaman/conditionalColoc.

## Introduction

Multimolecular interactions are the basis of cellular functions. High-resolution and super-resolution fluorescence microscopy allow us to routinely image three, four, or even more molecular entities at the same time within a cell (Huang et al., 2009; Sahl et al., 2017; Valm et al., 2017). Such multicolor imaging may provide a wealth of information about the spatiotemporal regulation of multimolecular interactions in their cellular context. However, without computational and quantitative analysis, much of this information remains unextracted (Bolte and Cordelières, 2006; Lagache et al., 2015).

Colocalization analysis is a cornerstone technique in cell biology for the analysis of multicolor microscopy images. It provides information on the localization of molecules within various subcellular compartments, shedding light on their function (Bolte and Cordelières, 2006; Dunn et al., 2011). For known molecular interactions, it allows their interrogation in their spatiotemporal cellular context, complementing biochemical and biophysical techniques, which detect interactions but taken out of their cellular context (Caetano et al., 2015; Helmuth et al., 2010; Lagache et al., 2018; Lagache et al., 2015). However, the overwhelming majority of colocalization analyses are applicable to only two-color microscopy images, i.e. they investigate only pairs of molecular entities. The few previous studies involving three-color colocalization were focused on either the question of whether the three imaged entities colocalize (Fletcher et al., 2010; Stauffer et al., 2018) or the question of mutually exclusive colocalization of one entity with the other two (Sastre et al., 2019). This greatly limits their applicability and the type of information that they may reveal, leading to underutilization of multicolor microscopy images.

Here we developed an approach for analyzing the colocalization relationships between three molecular entities, termed “conditional colocalization analysis,” based on spatial point pattern analysis of detected objects in microscopy images (Diggle, 2014; Helmuth et al., 2010; Lagache et al., 2015; Pastorek et al., 2016). Going beyond the question of whether colocalization is present or not, it addresses the question of whether the colocalization between two molecular entities is influenced, positively or negatively, by their colocalization with a third entity. For molecules known to interact (as identified via biochemical and/or biophysical techniques), the extracted colocalization relationships provide evidence for the cooperativity or competition between interactions, and for the enhancement or suppression of interactions at particular subcellular locations, domains or compartments.

To highlight the capabilities of conditional colocalization analysis, we applied it to two problems addressing different facets of molecular interactions and their regulation. In the first, we investigated the colocalization relationships between a receptor and its downstream effector in the context of functionally relevant plasma membrane locations, shedding light on the spatial compartmentalization of these interactions. In the second, we investigated the hierarchy of molecular interactions between three proteins, as a first step toward understanding the network of molecular interactions in which these molecules are involved.

## Results

### Derivation of conditional colocalization measures

To derive measures that reflect the extent to which one molecular entity influences – positively or negatively – the colocalization between two other molecular entities, we took a spatial point pattern analysis approach (Diggle, 2014; Helmuth et al., 2010; Lagache et al., 2015; Pastorek et al., 2016). Specifically, objects (i.e. the labeled molecular entities) in the images were first detected or segmented, and then their colocalization was assessed based on their nearest neighbor distance. We took this approach for two main reasons. First, for a basic measure of the extent of colocalization between two object types, the nearest neighbor distance allowed us to calculate the fraction of one object type colocalized with another object type (Helmuth et al., 2010; Lachmanovich et al., 2003). By interpreting colocalization fractions as probabilities, we were able to extend the analysis to calculate conditional probabilities and similar measures that reflected the relationships between object colocalizations, thus allowing us to address the question of the influence of one molecular entity on the colocalization of two other molecular entities. Second, object-based analysis offered more robust means to computationally assess the significance of any observed colocalization than pixel-based analysis such as the Pearson correlation function, as it allowed for (computationally) abolishing the relationships between objects (e.g. by object randomization) while maintaining individual object integrity (Fletcher et al., 2010; Lagache et al., 2015).

Throughout this work, we will employ the following terminology to refer to the three object types the colocalization relationships of which are being analyzed. The objects will be referred to as target (T), reference (R) and condition (C). Using this terminology, the question that the analysis aims to answer is: *How much is the colocalization of target objects with reference objects influenced by target and/or reference colocalization (or not) with condition objects?* To answer this question, we employed a four-step analysis process. For simplicity, the below descriptions and derivations are for images containing diffraction-limited, punctate objects, the positions of which are described by their center coordinates (determined, with sub-pixel precision, by fitting the signal with a Gaussian that approximates the microscope point spread function (Aguet et al., 2013; Jaqaman et al., 2008)). In the next section, we present an extension to the case when one channel contains non-punctate objects.

#### Step 1. Divide target and reference into groups colocalized or not colocalized with condition

Suppose there are *N_T_*, *N_R_* and *N_C_* target, reference and condition objects, respectively (**Figure 1A**). First, we determined for each target object and each reference object whether it colocalized with a condition object (**Figure 1A, B**). Those that colocalized with a condition object were in the condition-positive groups (*TwC* and *RwC*; “w” indicates “with”), while those that did not colocalize with any condition object were in the condition-negative groups (*TnC* and *RnC*; “n” indicates “without”).

**Figure 1.**
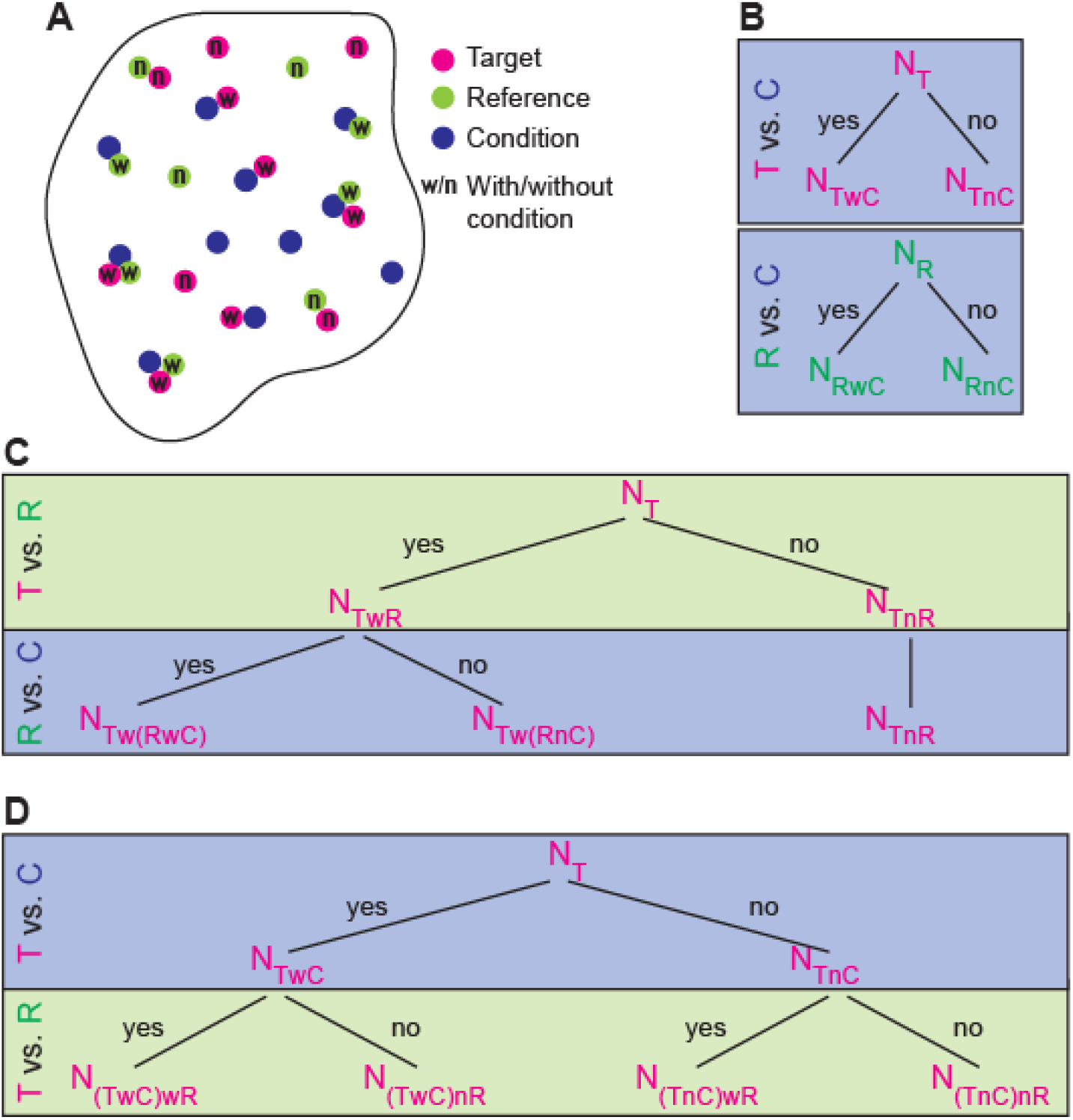
Schematic and decision trees for classifying target and reference objects in terms of their colocalization with condition objects and with each other. **(A)** Schematic of a “cell region” showing target (magenta) and reference (green) objects classified as with (w) or not with (n) condition objects (blue). **(B)** Decision trees for dividing target (top) and reference (bottom) objects into two groups each based on their colocalization (“yes” branch) or not (“no” branch) with condition objects. **(C)** Decision tree for first dividing target objects into colocalized or not with reference objects (top) and then, for target colocalized with reference, dividing them further based on reference colocalization with condition (bottom). **(D)** Decision tree for first dividing target objects into colocalized or not with condition (top) and then dividing each subgroup further based on target colocalization with reference.

Two objects were considered colocalized when the distance between their centers was below a particular colocalization radius. The colocalization radius could be set differently for each pair of channels, to account for the objects’ localization precision and the registration shift between each pair of channels. An important point to remember regarding localization precision is that, by definition, one detects the fluorophore labeling the object of interest, and not the object of interest itself. Thus, the true object localization precision will most likely be lower than the localization precision reflecting the signal-to-noise ratio (SNR) of the images (Bates et al., 2007).

This division yielded the probability (*p*) of target and reference objects to colocalize with condition objects:

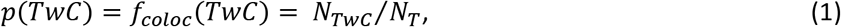

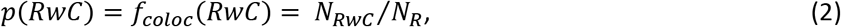

where *N_TwC_* and *N_RwC_* are, respectively, the number of target objects and reference objects colocalized with condition objects. In **Eqs. 1, 2** (and all throughout this work), *p*(…) and *f_coloc_*(…) indicate colocalization probability and colocalization fraction, respectively. These colocalization probabilities are equivalent to the colocalization measure proposed in (Helmuth et al., 2010) (where they are called “*C^T^*“).

#### Step 2. Calculate the fraction of target colocalizing with reference for the different group combinations

Next, we calculated the fractions of target objects in the different groups colocalized with reference objects in the different groups, also using the colocalization algorithm of (Helmuth et al., 2010). As in Step 1, two objects were considered colocalized when the distance between their centers was below a particular colocalization radius. From this, we obtained five colocalization fractions: *f_coloc_*(*TwR*) (for all target and reference), *f_coloc_*((*TwC*)*wR*) (for the condition-positive group of target), *f_coloc_*(*TnC*(*TwR*) (for the condition-negative group of target), *f_coloc_*(*Tw*(*RwC*)) (for the condition-positive group of reference), and *f_coloc_*(*Tw*(*RnC*)) (for the condition-negative group of reference).

For each colocalization fraction, we also calculated a corresponding coincidental colocalization fraction in the absence of any true relationship between target and reference 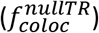. For this, we kept the reference objects in place, and replaced the target objects with points on a grid within the cell area, as described in (Helmuth et al., 2010). Using the same colocalization radius as for the original data, we calculated 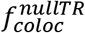 corresponding to each of the colocalization fractions mentioned above. A strength of this procedure is that it calculates the expected colocalization given the number and spatial distribution of the reference objects, i.e. it considers the spatial context of the colocalization analysis (Helmuth et al., 2010).

#### Step 3. Calculate conditional colocalization measures from colocalization fractions

The colocalization fractions obtained in Step 2 yielded the following conditional colocalization measures:

- **The overall colocalization probability (*p*(*TwR*)):** The all target vs. all reference analysis (**Figure 1C**, top row) yielded the overall probability of target to colocalize with reference, regardless of either’s relationship to condition:

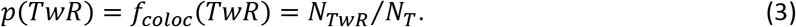 *N_TwR_* is the number of target objects colocalized with reference objects, irrespective of either’s relationship to condition objects (**Figure 1C**, top row).
- **Influence of target colocalization with condition on target colocalization with reference (*p*(*TwR*|*TwC*) and *p*(*TwR*|*TnC*)):** For this, the condition-positive target vs. all reference and condition-negative target vs. all reference analyses (**Figure 1D**) yielded:

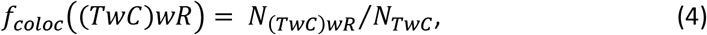

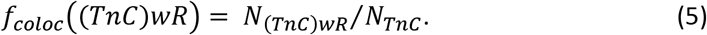 *N_(TwC)wR_* is the number of condition-positive target objects colocalized with reference objects (i.e. target objects colocalized with both reference objects and condition objects) (**Figure 1D**, left branch). *N_(TnC)wR_* is the number of condition-negative target objects colocalized with reference objects (i.e. target objects colocalized with reference objects but not with condition objects) (**Figure 1D**, right branch). *N_TwC_* and *N_TnC_* are the number of condition-positive and condition-negative target objects, respectively (from Step 1). It follows from **Eqs. 4, 5** that *f_coloc_*((*TwC*)*wR*) and *f_coloc_*((*TnC*)*wR*) indicate the conditional probabilities of target to colocalize with reference given that target is, respectively, colocalized, or not colocalized, with condition:

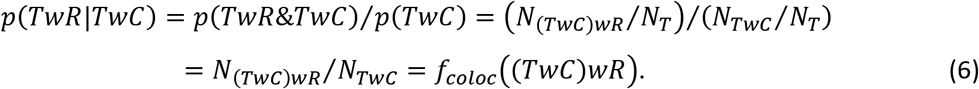

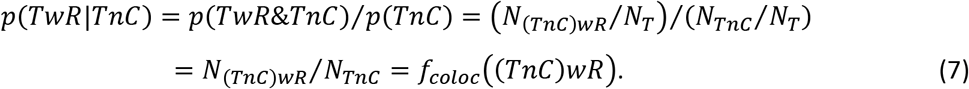 These are useful measures for conditional colocalization analysis, as their comparison to each other and to *p*(*TwR*) would answer the question whether the extent to which target colocalization with reference is influenced by target colocalization (or not) with condition.
- **Influence of reference colocalization with condition on target colocalization with reference (*p^rs^*(*Tw*(*RwC*)) and *p^rs^*(*Tw*(*RnC*))):** For this, the all target vs. condition-positive reference and all target vs. condition-negative reference analyses (**Figure 1C**, bottom row) yielded:

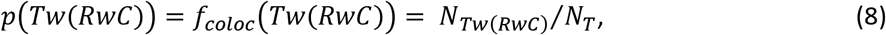

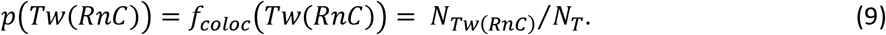 *N_Tw(Rwc)_* is the number of target objects colocalized with reference objects that are themselves colocalized with condition objects. *N_Tw(RnC)_* is the number of target objects colocalized with reference objects that are not colocalized with condition objects (**Figure 1C**, bottom row). While this analysis yielded the probabilities of target to colocalize with the indicated subsets of reference objects, these probabilities were not useful as is for answering the question whether the extent of target colocalization with reference is influenced by reference colocalization (or not) with condition. The reason is that the number of reference objects used in the different analyses is different. As a result, *N_RwC_* + *N_RnC_* = *N_R_* (**Figure 1B**, bottom), *N_Tw(RwC)_* + *N_Tw(RnC)_* = *N_TwR_* (**Figure 1C**, bottom row), and *p*(*Tw*(*RwC*)) + *p*(*Tw*(*RnC*)) = *p*(*TwR*) (**Eqs. 8, 9** vs. **Eq. 3**). However, in the absence of any special relationship between target objects and condition-positive or condition-negative reference objects, *p*(*Tw*(*RwC*)) is expected to be *p*(*TwR*) multiplied by *N_RwC_/N_R_* (= *p*(*RwC*); **Eq. 2**), while *p*(*Tw*(*RnC*)) is expected to be *p*(*TwR*) multiplied by *N_RnC_/N_R_* (= *p*(*RnC*) = 1 – *p*(*RwC*)). Thus, for a measure of the influence of reference colocalization with condition on the colocalization of target with reference, we rescaled the colocalization probabilities to factor out *p*(*RwC*) and *p*(*RnC*).

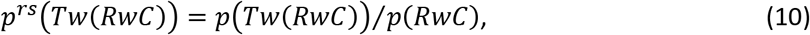

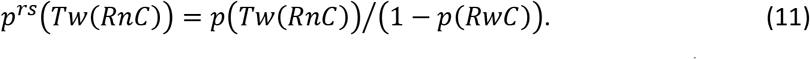 Note that these measures could be > 1, and thus they no longer indicated probabilities (unlike the original fractions in **Eqs. 8, 9**). Nevertheless, the rescaling factored out the number of reference objects used for analysis, allowing us to compare *p^rs^*(*Tw*(*RwC*)) and *p^rs^*(*Tw*(*RnC*)) to *p*(*TwR*), thus making these rescaled probabilities useful measures for assessing the extent to which target colocalization with reference is influenced by reference colocalization (or not) with condition. In summary, **Eqs. 3, 6, 7, 10 and 11** provided us with useful measures for conditional colocalization analysis, to answer the question whether the extent of target colocalization with reference is influenced by either’s colocalization with the condition. Using 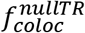 instead of *f_coloc_* in these equations yielded the coincidental counterparts of these measures in the absence of any true relationship between target and reference (*p_nullTR_* or 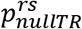).

#### Step 4. Assess significance of condition’s influence by randomizing condition locations

A critical issue for any colocalization analysis is to determine whether the observed colocalization is significantly different from what is expected by chance (Bolte and Cordelières, 2006; Fletcher et al., 2010; Lagache et al., 2015). The procedure in Steps 2 and 3 for calculating 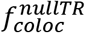 allowed us to compare the colocalization measures obtained for the data to their coincidental counterparts in the absence of a true relationship between target and reference. However, a similar concern applied to the extracted influence of the condition on target colocalization with reference, and this was not covered by calculating 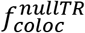.

To address this concern, we randomized the locations of the condition objects, while keeping the same number of condition objects and not altering the target or reference objects in any way (see Materials and Methods and **Suppl. Figure S1**). We then repeated Steps 1-3 above. Comparison of the original data colocalization measures to those calculated after condition location randomization (*p_randC_* or 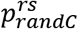) allowed us to determine whether colocalization with the condition significantly increased (or decreased) the extent of target colocalization with reference.

### Generalization of conditional colocalization analysis to non-punctate objects

In the above analysis scheme, we assumed that the objects were diffraction-limited punctate objects and defined the distance between objects as the distance between their centers. While this covers a large range of colocalization problems, including of molecules and of sub-resolution subcellular structures such as clathrin-coated pits and other nanodomains (Aguet et al., 2013; Sezgin et al., 2017), many subcellular structures of interest, especially as condition objects, could be larger than the resolution limit.

Therefore, we generalized the above scheme to the case where one out of the three object types was non-punctate. For this, we introduced two modifications: (i) We replaced Gaussian fitting with image segmentation to delineate objects in the non-punctate channel. (ii) We defined the distance between a punctate object and a non-punctate object as the distance from the center of the punctate object to the nearest pixel of the non-punctate object.

With this, if the non-punctate objects were the condition, the above derivations and calculations were applicable as is and they yielded all five conditional colocalization measures of interest. If the non-punctate objects were the reference, the above analysis scheme allowed us to calculate three out of the five measures: *p*(*TwR*), *p*(*TwR|TwC*) and *p*(*TwR|TnC*). We did not calculate the remaining measures because they would require classifying larger, non-punctate reference objects as colocalized or not with smaller, punctate condition objects, which is not clearly defined. For the same reason, the above analysis was not generalized to non-punctate target objects, which would have to be classified as colocalized or not with both punctate reference objects and punctate condition objects, both of which would be smaller than the target objects in this scenario.

### Analysis scheme accurately estimates conditional colocalization measures in simulated data

To assess the accuracy of our conditional colocalization analysis scheme, we tested it on simulated data with known ground-truth conditional colocalization measures. To make the simulations as realistic as possible, we took experimental data cell masks and simulated various colocalization scenarios within the cell masks. To localize punctate objects, we selected sub-pixel coordinates within each cell mask. For non-punctate objects, we used paxillin patch segmentations from our experimental data (belonging to the cell masks employed). This allowed us to simulate non-punctate objects of varying sizes and shapes in a realistic manner, especially in the context of our biological applications. Further simulation details are described in Materials and Methods.

First, we validated the core colocalization analysis between two sets of objects. For the case of punctate target and reference objects, we used *N_T_* = 150, 200, 250, 300, 350, *N_R_* = 150, 200, 250, 300, 350 and *p*(*TwR*) = 0, 0.25, 0.5, 0.75, 1 (i.e. 125 parameter combinations). We generated 28 instances of each of these parameter combinations, each in a different cell mask. Using a colocalization radius of three pixels (as used for our experimental data analysis, as described below), the calculated *p*(*TwR*) for each parameter combination was very close to its ground-truth value, and, as expected, it was independent of *N_T_* and *N_R_* (**Figure 2A**, left). Also as expected, the coincidental colocalization probability *p_nullTR_*(*TwR*) (calculated by replacing target object positions with a grid (Helmuth et al., 2010)) depended only on *N_R_*, reflecting the fraction of the cell mask area covered by the colocalization zones around the reference objects (**Figure 2A**, right).

**Figure 2.**
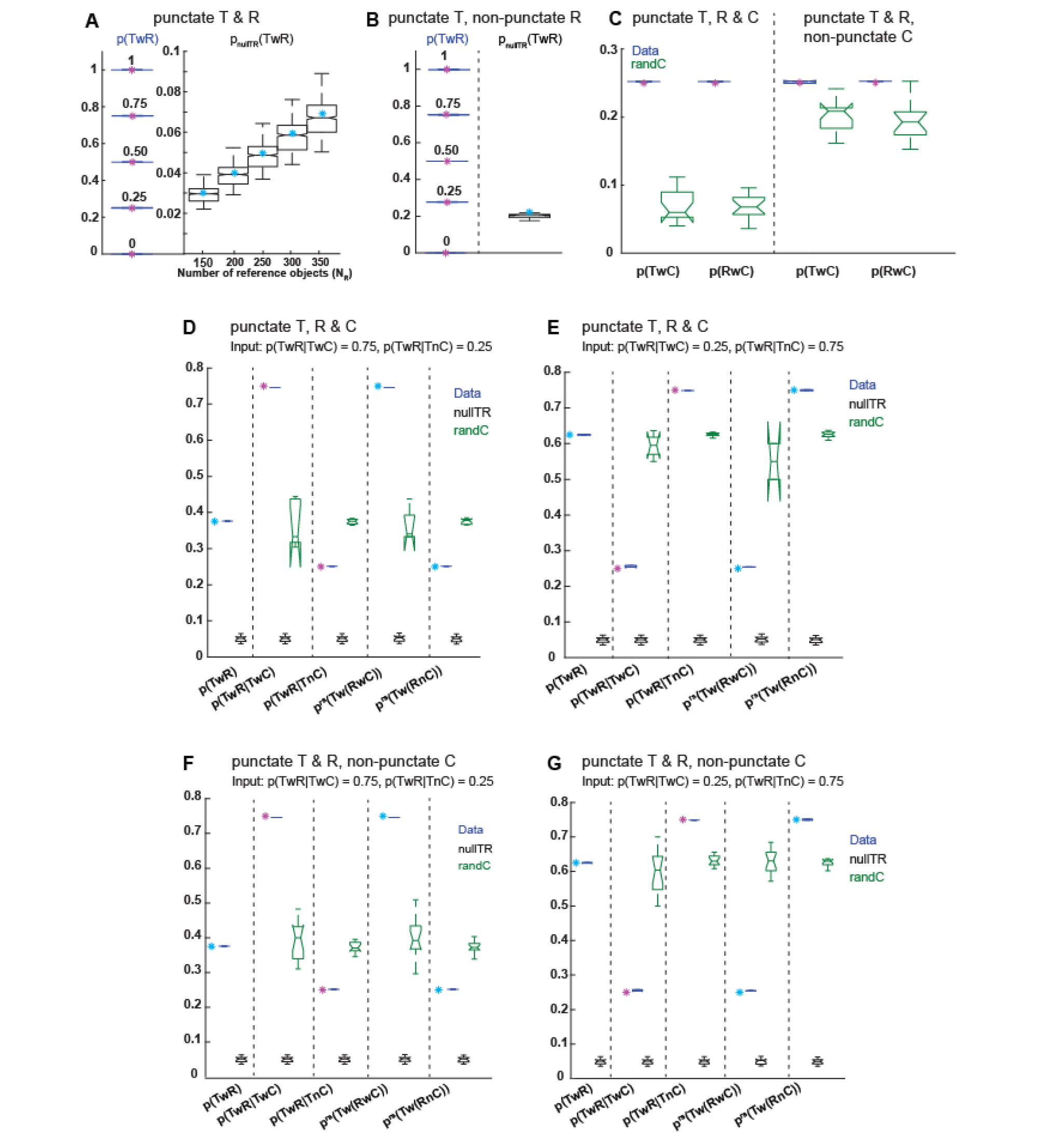
Analysis scheme accurately estimates conditional colocalization measures. **(A, B)** Calculated *p*(*TwR*) (left) and corresponding *p_nullTR_*(*TwR*) (right) for simulations of punctate **(A)** and non-punctate **(B)** reference with punctate target. Each box shows the calculated values for all parameter combinations using the same input *p*(*TwR*) or having the same expected *P_nullTR_*(*TwR*). Input/expected values are shown as magenta/cyan asterisks. For each box, the central mark is the median, the edges are the 25th and 75th percentiles, and the dashed whiskers extend to the most extreme data points. Notch around median indicates the 95% confidence interval of the median. In (A), each box represents N = 700 values (25 *N_T_* and *N_R_* combinations, or 25 *N_T_* and *p*(*TwR*) combinations, each repeated 28 times). In (B), each *p*(*TwR*) box represents N = 140 values (5 *N_T_* values, each repeated 28 times), while the *p_nullTR_*(*TwR*) box represents N = 700 values (25 *N_T_* and *p*(*TwR*) combinations, each repeated 28 times). **(C-G)** Representative results of calculated conditional colocalization measures. Each box represents N = 15 repeats of the stated simulation parameters. **(C)** Calculated *p*(*TwC*) and *p*(*RwC*) and their “randC” counterparts for simulations of punctate (left) and non-punctate (right) condition with punctate target and reference. Simulation parameters: *p*(*TwR*|*TwC*)=0.75, *p*(*TwR*|*TnC*)=0.25, *p*(*TwC*)=*p*(*RwC*)=0.25. Input values are shown as magenta asterisks. **(D-G)** Calculated conditional colocalization measures and their “nullTR” and “randC” counterparts for simulations of punctate **(D, E)** and non-punctate **(F, G)** condition with punctate target and reference, using the indicated *p*(*TwR*|*TwC*) and *p*(*TwR*|*TnC*) as well as *p*(*TwC*)=*p*(*RwC*)=0.25. Input/expected values are shown as magenta/cyan asterisks (see Materials and Methods for details).

For the case of punctate target objects and non-punctate reference objects, we varied *N_T_* = 150, 200, 250, 300, 350 and *p*(*TwR*) = 0, 0.25, 0.5, 0.75, 1, while taking the reference objects from our experimental data paxillin patch segmentations (i.e. 25 parameter combinations in this case). For this case too, the calculated *p*(*TwR*) for each parameter combination was very close to its ground-truth value, and *P_nullTR_*(*TwR*) reflected the fraction of the cell area covered by the reference objects and the colocalization zones around them (**Figure 2B**). These tests indicate that the core colocalization analysis between two sets of objects yields accurate results for both the case of two sets of punctate objects and the case of punctate target objects and non-punctate reference objects.

Next, we validated the conditional colocalization analysis. Here, given our observation that the estimated *p*(*TwR*) is equally accurate for all tested *N_T_* and *N_R_* (**Figure 2A, B**), we fixed *N_T_* and *N_R_* at 250 (middle of the range tested above) and only varied *N_C_* and the various colocalization probabilities. For the case of all punctuate objects, we varied *N_c_* = 250, 300 and 350, *p*(*TwC*) = 0.25, 0.5, 0.75, *p*(*RwC*) = 0.25, 0.5, 0.75, *p*(*TwR|TwC*) = 0.25, 0.5, 0.75 and *p*(*TwR|TnC*) = 0.25, 0.5, 0.75 (i.e. 243 parameter combinations in total). For the case of non-punctate condition objects, we took the condition objects from our experimental data paxillin patch segmentations (as above, but now as condition objects instead of reference objects), and varied *p*(*TwC*), *p*(*RwC*), *p*(*TwR|TwC*) and *p*(*TwR|TnC*) as above (i.e. 81 parameter combinations). Each of these parameter combinations was simulated 15 times using 15 cell masks (and their associated paxillin patch segmentations in the case of non-punctate condition objects). For each parameter combination, we compared the calculated *p*(*TwC*), *p*(*RwC*), *p*(*TwR|TwC*) and *p*(*TwR|TnC*) to their input values, and the calculated *p*(*TwR*), *p^rs^*(*Tw*(*RwC*)) and *p^rs^*(*Tw*(*RnC*)) to their expected values (see **Eqs. S1, S4 and S5** in Materials and Methods). We found that all conditional colocalization measures were estimated accurately, regardless of number and type (punctate vs. non-punctate) of condition objects (**Figure 2C-G**). The equivalent results for punctate and non-punctate condition objects imply that our validation test results are not limited to paxillin patch-like condition objects, as used in these validation simulations, but can be generalized to condition objects of arbitrary shape and size.

### Specificity: analysis scheme correctly identifies no colocalization in an experimental negative control

While the above simulations verified that our analysis scheme was able to accurately estimate the conditional colocalization measures, simulations might not capture some of the complexity and nuances of cellular images. For example, in our simulations, objects were placed randomly within the cell mask. However, molecules in a cell are often distributed in an inhomogeneous manner, leading to nonspecific colocalization that is not related to any direct or indirect interactions. Furthermore, the resolution limit of light microscopy, and the colocalization radius used for the analysis of diffraction-limited images, are relatively large when compared to molecular dimensions. In our analyses, we employed a colocalization radius of three pixels (equal to 243 nm or 267 nm depending on dataset/microscope), in order to account for registration shifts between different channels and to gather enough statistics per group of objects. Therefore, an important question to address was whether nonspecific colocalization might be detected in the absence of any true (direct or indirect) interactions and specific relationships between the molecules of interest.

To address this question, we applied conditional colocalization analysis to an experimental negative control dataset. We performed three-color immunofluorescence (IF) imaging of TfR (transferrin receptor; taken as “target”), TSAd (taken as “reference”) and CHC (clathrin heavy chain; taken as “condition”) in endothelial cells (ECs). We employed total internal reflection fluorescence microscopy (TIRFM), with a penetration depth of ~90 nm, to focus on the bottom cell surface and minimize apparent colocalization from signals existing in different z-planes (**Figure 3A**). TfR is a cell surface receptor responsible for binding transferrin, an iron-binding glycoprotein (Gammella et al., 2017). On the other hand, TSAd is an adaptor protein that, in ECs, binds to activated VEGFR-2 (Sun et al., 2012). There are no reports of association between TSAd and TfR, making this pair of molecules a good negative control. As for CHC, it is a marker for clathrin-coated structures (CCSs), including clathrin-coated pits, which are responsible for clathrin-mediated endocytosis, the main mechanism for TfR internalization into the cell (Liu et al., 2010; Mayle et al., 2012). Thus, we would expect a certain level of association between TfR and CCSs, making them relevant condition objects for conditional colocalization analysis.

**Figure 3:**
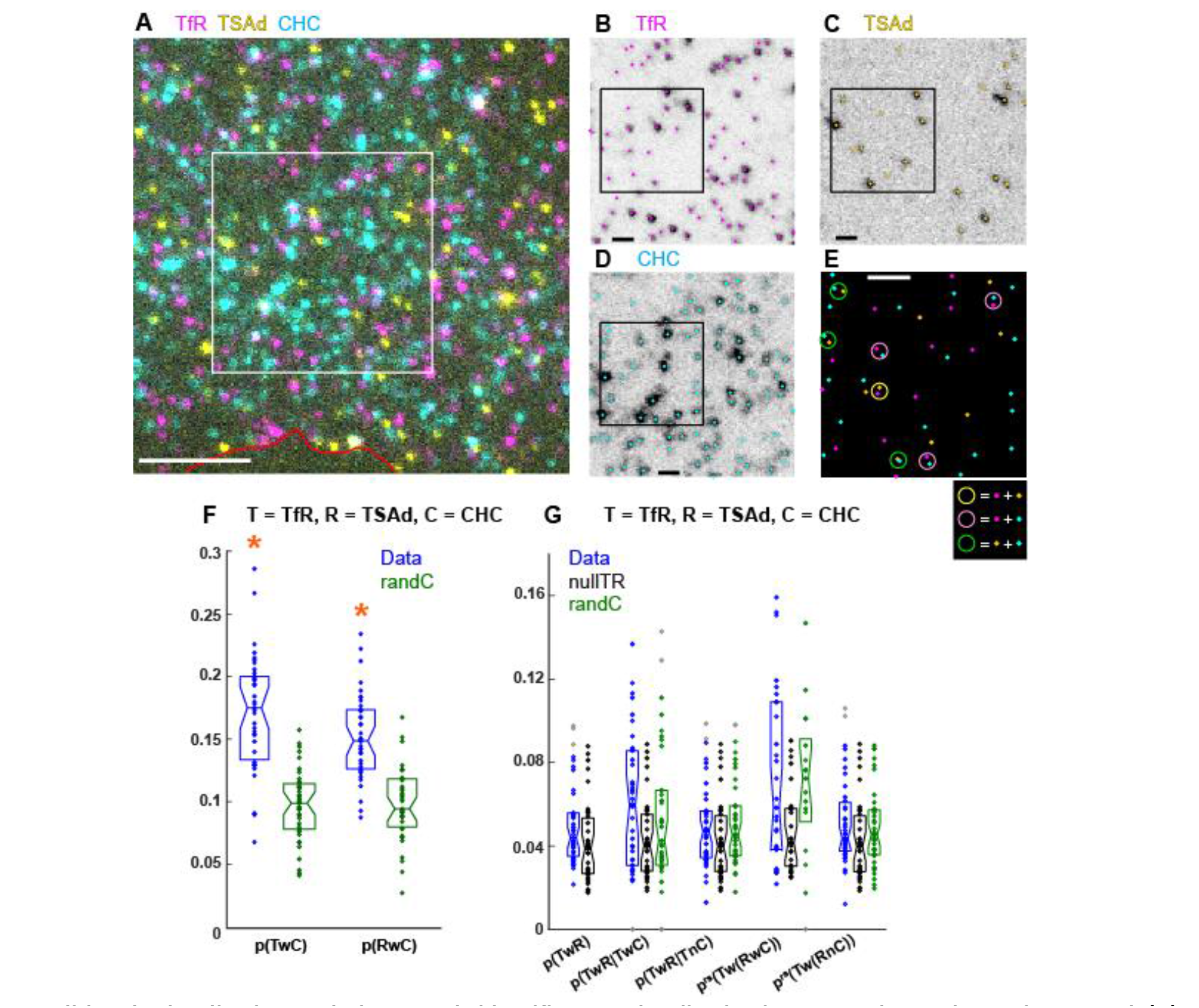
Conditional colocalization analysis correctly identifies no colocalization in an experimental negative control. **(A)** Representative three-color IF image of TfR, TSAd and CHC on the surface of a TIME cell imaged via TIRFM. Red line shows cell edge. Scale bar, 5 μm. **(B-D)** Particle detections (shown as dots) overlaid on the individual channels for the area within the white box in (A). Scale bar, 1 μm. **(E)** Overlay of the three channel detections for the area within the black box in (B-D), with colorcoding following that in (B-D). Colored circles (diameter = 243 nm) point out colocalization events between the different molecules, following color-coding in legend. Scale bar, 1 μm. **(F)** Probabilities of target (TfR) and reference (TSAd) to colocalize with condition (CHC), together with their coincidental counterparts (“randC”). For each box, individual dots indicate individ ual cell measurements, the central mark is the median and the edges are the 25th and 75th percentiles. Notch around median indicates the 95% confidence interval of the median. Dots with same color as box are inliers and gray dots are outliers (as deemed by the Matlab “boxplot” function). Orange asterisks (*): p < 0.01, indicating that the probability of colocalization with condition is significantly greater than its coincidental counterpart, as assessed via a Wilcoxon rank-sum test. **(G)** Conditional colocalization measures and their coincidental counterparts (“nullTR” and “randC”) for same object combination as in (F). Boxplots and dots as in (F). None of the conditional colocalization measures pass the significance test (See text), and thus none are highlighted. N = 39 cells from 4 repeats. See **Table 3** for number of objects per channel and other dataset properties.

All channels were of a punctate nature. Thus, we determined the object positions in each channel with subpixel localization using Gaussian fitting (Aguet et al., 2013) (**Figure 3B-E**), and then applied the conditional colocalization analysis algorithm for three sets of punctate objects. As expected, TfR exhibited significant colocalization with CCSs (the condition objects) (**Figure 3F**). TSAd also exhibited significant colocalization with CCSs, although less than TfR (**Figure 3F**). TSAd colocalization with CCSs was probably because of its interactions with cell surface receptors such as VEGFR-2, which themselves might reside in CCSs. Overall, both molecules colocalized with CCSs, giving us a sizable condition-positive group of each molecule type for conditional colocalization analysis.

Importantly, even though both TfR and TSAd colocalized with CCSs, TfR did not exhibit any significant colocalization with TSAd by any of our conditional colocalization measures (**Figure 3G**). No measure was statistically distinguishable from its corresponding coincidental counterparts (nullTR and randC (**Figure 3G**; blue vs. black vs. green)) and *p*(*TwR*). Overall, these negative control results demonstrate that conditional colocalization analysis possesses reliable specificity, correctly identifying situations of no colocalization. This lends confidence to the conditional colocalization measures and the colocalization relationships that they reveal.

### Sensitivity: analysis scheme captures the colocalization properties of an experimental positive control

In addition to the properties of cellular images that could undermine the specificity of conditional colocalization analysis, some factors might also undermine the sensitivity of the analysis, i.e. its ability to detect colocalization when the co-imaged molecules are known to interact. Such factors include antibody specificity and sensitivity in the case of IF and detection and segmentation errors related to image SNR (including false positives and false negatives). To test the sensitivity of our conditional colocalization analysis scheme, we applied it to images of molecules known to interact with each other.

For this, we chose three proteins that are part of focal adhesion (FA) complexes: β_1_-integrin, FAK (focal adhesion kinase) and paxillin (Geiger et al., 2009; Vicente-Manzanares and Horwitz, 2011). Integrins are a family of cell surface receptors that bind to components of the extracellular matrix and initiate the formation of FAs (Bachmann et al., 2019; Hynes, 2002). FA formation involves the activation of a series of kinases, such as FAK, which in turn activates paxillin, an FA adapter protein, responsible for the scaffolding and recruitment of other signaling molecules to FAs (López-Colomé et al., 2017; Mitra et al., 2005). Given their mutual interactions and involvement in FA formation, activated β_1_-integrin, activated FAK (phosphorylated FAK or pFAK), and paxillin constituted a viable positive control for our conditional colocalization analysis.

We performed three-color IF imaging of activated β_1_-integrin, pFAK, and paxillin in ECs using TIRFM (**Figure 4A-E**). For activated β_1_-integrin and pFAK, we used low antibody concentrations, and further bleached the images (see Materials and Methods), in order to obtain punctate signals, which were detected with sub-pixel localization using Gaussian fitting (Aguet et al., 2013) (**Figure 4B, C, E**). On the other hand, the paxillin signal was non-punctate, and it was segmented using intensity-based thresholding (Jaqaman et al., 2016), focusing on objects larger than the diffraction limit (**Figure 4D, E**). Thus, paxillin was used as a marker for FAs, which served as the condition for our analysis. Note that the incomplete labeling of activated β_1_-integrin and pFAK posed an additional challenge for conditional colocalization analysis, as incomplete labeling is expected to reduce the extent of observed colocalization between the imaged molecules.

**Figure 4:**
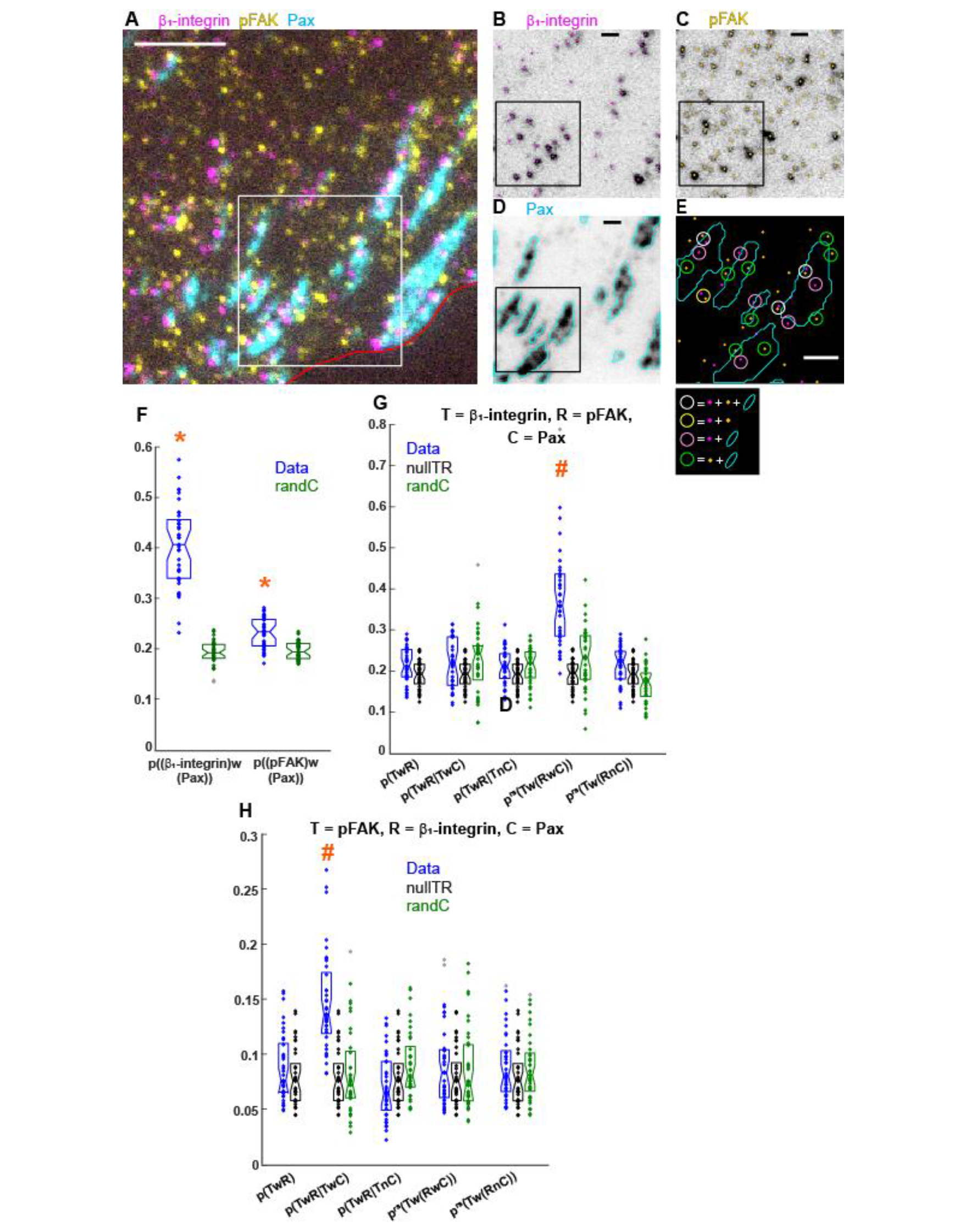
Conditional colocalization analysis captures the colocalization properties of an experimental positive control. **(A)** Representative three-color IF image of activated β1-integrin, pFAK and paxillin on the surface of a TIME cell imaged via TIRFM. Red line shows cell edge. Scale bar, 5 μm. **(B-D)** Particle detections (shown as dots in (B, C)) and patch segmentation (shown as outlines in (D)) overlaid on the individual channels for the area within the white box in (A). Scale bar, 1 μm. **(E)** Overlay of the three channel detections and segmentations for the area within the black box in (B-D), with color coding following that in (B-D). Colored circles (diameter = 243 nm) point out colocalization events between the different molecules, following color-coding in legend. Scale bar, 1 μm. **(F)** Probabilities of β_1_-integrin and pFAK to colocalize with the condition (paxillin; representing FAs), together with their coincidental counterparts (“randC”). Boxplots, dots and orange asterisks as in Figure 3F. **(G, H)** Conditional colocalization measures and their coincidental counterparts (“nullTR” and “randC”) for target = β_1_-integrin and reference = pFAK **(G)** and vice versa **(H)**, both with condition = paxillin. Boxplots and dots as in Figure 3F. Orange pound signs (#): colocalization measure passes the significance test, i.e. it is significantly greater than its coincidental counterparts (both “nullTR” and “randC”) and *p*(*TwR*), as assessed via three Wilcoxon rank-sum tests. The significance threshold for each individual test is calculated using the Dunn–Sidak correction to give a total type-I error rate (for all three tests) of 0.05. N = 37 cells from 4 repeats. See **Table 3** for number of objects per channel and other dataset properties.

Both activated β_1_-integrin and pFAK exhibited significant colocalization with the paxillin patches, with activated β_1_-integrin showing greater colocalization than pFAK (**Figure 4F**). They also exhibited significant colocalization with each other, but only for the subpopulation of pFAK that was colocalized with paxillin; this was revealed by performing the conditional colocalization analysis once taking β_1_-integrin as target and pFAK as reference (**Figure 4G**), and once taking pFAK as target and β_1_-integrin as reference (**Figure 4H**). With pFAK as reference, only *p^rs^*(*Tw*(*RwC*)) (RwC = pFAK with paxillin) was significantly greater than its coincidental counterparts and *p*(*TwR*) (**Figure 4G**). Conversely, with pFAK as target, only *p*(*TwR|TwC*) (TwC = pFAK with paxillin) was significantly greater than its coincidental counterparts and *p*(*TwR*) (**Figure 4H**). Of note, in neither case was the overall colocalization probability *p*(*TwR*) different from its coincidental counterpart. In other words, only by identifying the proper subset of pFAK – those colocalized with FAs – was it possible to capture the colocalization relationship between pFAK and β_1_-integrin. The pFAK molecules not colocalized with FAs were probably interacting with cell surface receptors other than integrins and involved in other signaling pathways (Marlowe et al., 2016; Murphy et al., 2019; Rigiracciolo et al., 2019; Sulzmaier et al., 2014).

These results highlight the power of conditional colocalization analysis in the face of molecular and interaction heterogeneity within the cell and demonstrate its ability to quantify the context-based colocalization relationships between (directly or indirectly) interacting molecules.

### Application 1: Conditional colocalization analysis reveals modulation of receptor-downstream effector colocalization at specific membrane locations

The spatial compartmentalization of molecular interactions is an emerging theme in signaling. Cell surface receptors have been observed to preferentially interact with downstream effectors at particular membrane microdomains, and to signal differently from different subcellular compartments (Birch et al., 2021; Delos Santos et al., 2015; Eichmann and Simons, 2012; Jaqaman and Ditlev, 2021; Sorkin and von Zastrow, 2009; Sungkaworn et al., 2017). Conditional colocalization analysis is ideally suited to shed light on the spatial regulation and/or compartmentalization of molecular interactions (and consequently signaling) within their cellular environment, in an automated and quantitative fashion.

To illustrate this capability, we applied conditional colocalization analysis to the receptor tyrosine kinase VEGFR-2 (Vascular Endothelial Growth Factor Receptor 2) and its downstream effector TSAd (T cell specific adaptor protein; also known as VEGF receptor adaptor protein (VRAP)) in the context of FAs. VEGFR-2, expressed on the surface of ECs, is the main receptor for VEGF (short for VEGF-A) (Ferrara et al., 2003; Simons et al., 2016). VEGFR-2 activation by VEGF triggers multiple signaling pathways, initiated by the binding of multiple downstream effectors to activated VEGFR-2 (Ferrara et al., 2003; Simons et al., 2016). Among the different downstream effectors, TSAd initiates a signaling cascade that modulates FA turnover and cell adhesion, critically important for EC motility and angiogenic sprouting (Abedi and Zachary, 1997; Birukova et al., 2009; Gordon et al., 2016; Sun et al., 2012; Wu et al., 2000). Motivated by our initial observation that VEGFR-2 tends to localize near FAs (**Figure 5A**), and given that TSAd binding to VEGFR-2 results in signaling to FAs, we employed conditional colocalization analysis to investigate whether the interactions between VEGFR-2 and TSAd are enhanced at FAs.

**Figure 5:**
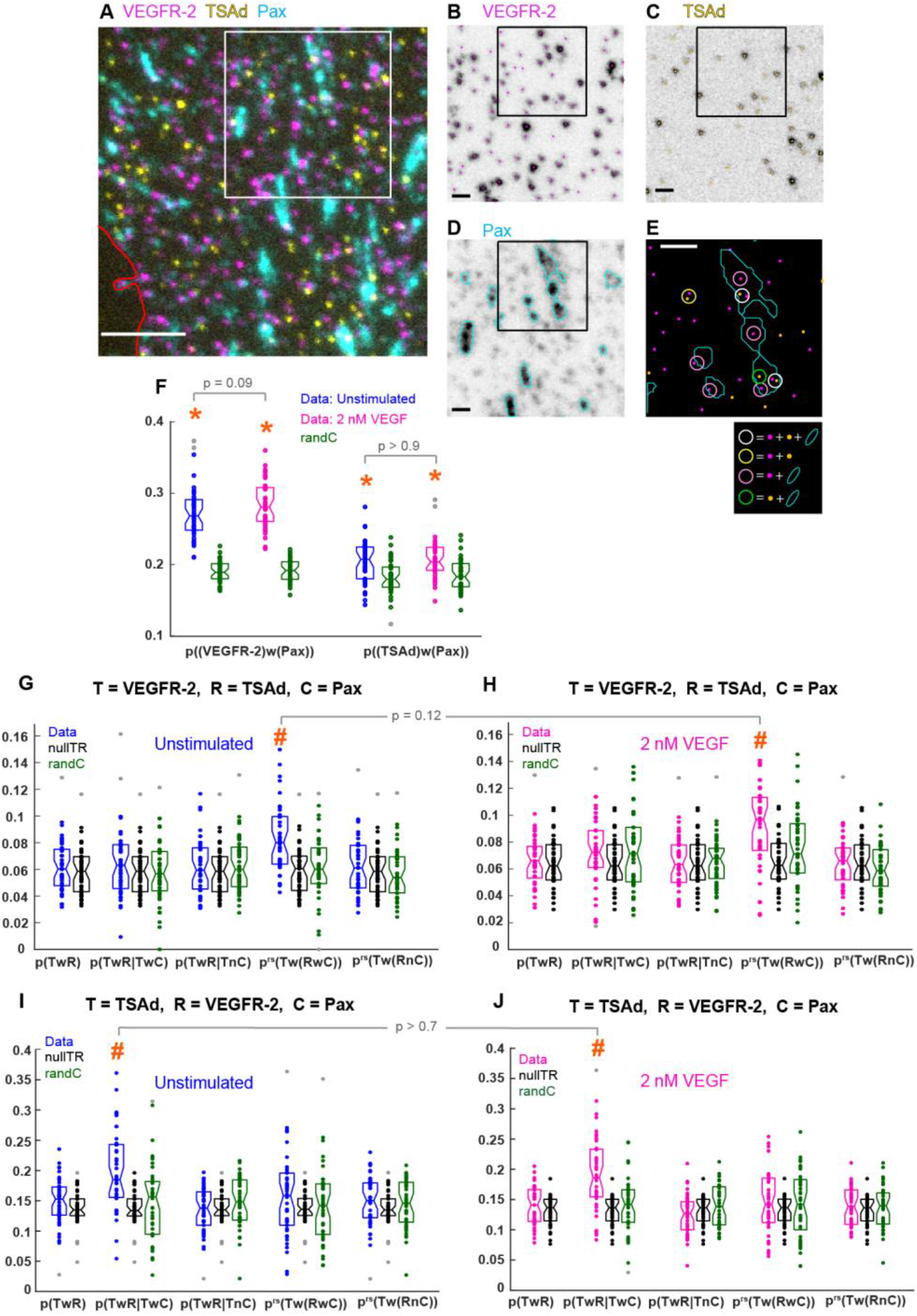
VEGFR-2-TSAd colocalization is enhanced for TSAd that is colocalized with FAs. **(A)** Representative three-color IF image of VEGFR-2, TSAd and paxillin on the surface of a TIME cell imaged via TIRFM. Red line shows cell edge. Scale bar, 5 μm. **(B-D)** Particle detections (shown as dots in (B, C)) and patch segmentation (shown as outlines in (D)) overlaid on the individual channels for the area within the black box in (A). Scale bar, 1 μm. **(E)** Overlay of the three channel detections and segmentations for the area within the black box in (B-D), with color coding following that in (B-D). Colored circles (diameter = 243 nm) point out the colocalization events between the different molecules, following color-coding in legend. Scale bar, 1 μm. **(F)** Probabilities of VEGFR-2 and TSAd to colocalize with the condition (paxillin; representing FAs), in the presence or absence of VEGF, together with their coincidental counterparts (“randC”). Boxplots, dots and orange asterisks as in Figure 3F. The probabilities in the absence and presence of VEGF were compared to each other using a Wilcoxon rank-sum test, resulting in indicated p-values. **(GJ)** Conditional colocalization measures and their coincidental counterparts (“nullTR” and “randC”) for target = VEGFR-2 and reference = TSAd **(G, H)** and vice versa **(I, J)**, both with condition = paxillin, in the absence **(G, I)** or presence **(H, J)** of 2 nM VEGF. Boxplots, dots and orange pound signs as in Figure 4G. *p^rs^*(*Tw*(*RwC*)|*TwC*) and *p^rs^*(*Tw(RnC)|TnC*) were omitted because they did not provide useful information and, at the same time, they threw off the y-axis range (as observed for the negative and positive control analyses in Figures 3, 4). Significant measures were compared between the absence and presence of VEGF as in (F), yielding indicated p-values. N = 41 cells from 4 repeats. See **Table 3** for number of objects per channel and other dataset properties.

We performed three-color IF imaging of VEGFR-2, TSAd and paxillin using TIRFM, in the absence or presence of 2 nM VEGF (5 min stimulation before fixation, which robustly activates VEGFR-2 (da Rocha-Azevedo et al., 2020)). The VEGFR-2 and TSAd signals were punctate and were thus localized using Gaussian fitting (Aguet et al., 2013) (**Figure 5B, C**). FAs, as marked by paxillin patches, were segmented using intensity-based thresholding as above (Jaqaman et al., 2016) (**Figure 5D**). Taking FAs as the condition, VEGFR-2 exhibited significant colocalization with FAs, both in the absence and presence of VEGF (**Figure 5E, F**). There was a mild, marginally significant increase in VEGFR-2 colocalization with FAs upon stimulation by VEGF (**Figure 5F**). TSAd also exhibited significant colocalization with FAs in the absence and presence of VEGF, although to a lesser extent than VEGFR-2 (**Figure 5F**).

For conditional colocalization analysis, taking VEGFR-2 as target and TSAd as reference, we found that VEGFR-2 exhibited significant colocalization with TSAd for the subset of TSAd colocalized with FAs (*p^rs^*(*Tw*(*RwC*)) in **Figure 5G, H**). This was the case both in the absence and presence of VEGF, with a mild (albeit not statistically significant) increase in colocalization in the presence of VEGF (**Figure 5G, H**). Performing the converse analysis, i.e. taking TSAd as target and VEGFR-2 as reference, maintained the special status of TSAd that is colocalized with FAs: here too, TSAd exhibited significant colocalization with VEGFR-2 only when TSAd colocalized with FAs (*p*(*TwR|TwC*) in **Figure 5I, J**). Our observation that the overall colocalization between VEGFR-2 and TSAd (*p*(*TwR*) in **Figure 5G-J**) was not significant under any circumstances highlights the need for conditional colocalization analysis and the unique information that it reveals, which would be lost with traditional colocalization analysis and other ensemble/bulk approaches.

To determine whether the effect of FAs on VEGFR-2 and TSAd colocalization was specific to FAs, we performed conditional colocalization analysis of VEGFR-2 and TSAd in the context of clathrin-coated structures (CCSs) and compared the analysis results to those in the context of FAs. VEGFR-2 is internalized via clathrin-mediated endocytosis in both ligand-dependent and ligand-independent manners (Basagiannis et al., 2016; Lampugnani et al., 2006). VEGFR-2 can also continue downstream signaling after internalization (Lanahan et al., 2010; Simons et al., 2016). These properties made CCSs a viable condition object with which we would expect VEGFR-2 to colocalize, possibly together with downstream effectors.

We performed three-color IF imaging of VEGFR-2, TSAd and CHC using TIRFM, in the absence and presence of VEGF as above. As all channels were punctate, all objects were localized using Gaussian fitting (Aguet et al., 2013) (**Suppl. Figure S2A-E**). Both VEGFR-2 and TSAd colocalized with CCSs as the condition (**Suppl. Figure S2F**). We performed conditional colocalization analysis taking VEGFR-2 as target and TSAd as reference and vice versa (**Suppl. Figure S2G-J**). No colocalization measures were significant, except for the probability of VEGFR-2 (target) to colocalize with TSAd (reference) if VEGFR-2 was colocalized with CCSs (condition), in the presence of VEGF (*p*(*TwR|TwC*) in **Suppl. Figure S2H**). Thus, overall, CCSs exert much less influence on the colocalization between VEGFR-2 and TSAd than FAs. Furthermore, their influence is different from that of FAs: In the case of FAs, colocalization was enhanced for the subset of TSAd at FAs. In the case of CCSs, colocalization was enhanced for the subset of VEGFR-2 at CCSs.

Altogether, these analyses indicate that the colocalization relationship between VEGFR-2 and TSAd varies depending on their location within the plane of the plasma membrane. In particular, it is influenced in a specific, positive manner by the colocalization of TSAd with FAs, which are functionally relevant for VEGFR-2 signaling downstream of TSAd. To the extent that colocalization is reflective of interactions between VEGFR-2 and TSAd, this raises the question of what is special about the subset of TSAd at FAs, such that VEGFR-2 exhibits increased colocalization with it. One possibility is the presence of other interaction partners or feedback from downstream signaling that stabilize the interactions between TSAd and VEGFR-2. Performing further conditional colocalization analysis with other interaction partners and combining it with perturbation of the different molecular players is expected to help determine the molecular mechanisms behind this increase in VEGFR-2-TSAd interactions at FAs. Importantly, the increased interactions most likely facilitate VEGFR-2 signaling in the right place at the right time to regulate cellular adhesion and migration in response to VEGF.

### Application 2: Conditional colocalization analysis reveals colocalization interdependencies between three molecules

In the cell, molecules are engaged in networks of interactions with many other molecules, leading to the formation of complexes, clusters and condensates that are important for cell signaling and other cellular functions (Gingras et al., 2019; Jaqaman and Ditlev, 2021; Yanez-Mo et al., 2009). Determining which molecules interact directly, and which interact indirectly through mediator molecules, is a critical step toward understanding the molecular mechanisms underlying the formation of these higher-order molecular assemblies. Through its analysis of the colocalization relationships between multiple molecules, conditional colocalization analysis is ideally suited to shed light on the hierarchy of intermolecular interactions within their cellular context.

This application of conditional colocalization analysis differs in purpose from the previous application. In the previous application, the condition objects were pre-determined subcellular structures/membrane microdomains (e.g. FAs or CCSs), and the question was whether these domains (the condition) played any role in the colocalization of two molecules. In this application, the goal is to identify the condition itself, i.e. the molecule that may act as a mediator between the other two.

To illustrate this capability, we applied conditional colocalization analysis to the three cell surface proteins CD36, β_1_-integrin and CD151. CD36 is a scavenger receptor expressed on the surface of many cell types, where it binds diverse ligands and is implicated in various (patho-)physiological processes, such as angiogenesis, atherosclerosis, Alzheimer’s disease and immunity (Silverstein and Febbraio, 2009). CD36 clustering and engagement within multimolecular complexes containing integrins (particularly those containing the β_1_ subunit) and tetraspanins (CD9, CD81 and/or CD151, dependent on cell type) are thought to be important for its signaling function, as the two short intracellular domains of CD36 lack any known signaling motifs (Githaka et al., 2016; Heit et al., 2013; Huang et al., 2011; Kazerounian et al., 2011; Miao et al., 2001; Primo et al., 2005; Thorne et al., 2000). However, these integrins and tetraspanins interact with each other and with other membrane proteins (Charrin et al., 2009; Yauch et al., 2000; Zhang et al., 2009), making it difficult to determine the molecular mechanisms underlying their interactions with CD36 and their contribution to CD36 signaling.

We reasoned that conditional colocalization analysis could shed light on the hierarchy, if any, in the interactions between CD36, integrins and tetraspanins. In ECs, β1-integrin and the tetraspanin CD151 have been shown to co-immunoprecipitate with CD36 (Kazerounian et al., 2011; Primo et al., 2005). Thus, we asked whether CD36 interacts separately (or independently) with β1-integrin and CD151, or whether its interactions with one depend on the other. Addressing this question with traditional knockout or knockdown approaches would not be straightforward, given the essential role that integrins play in cell physiology and the redundancy between tetraspanins (Charrin et al., 2009; Winograd-Katz et al., 2014).

We performed fixed cell imaging of CD36, β1-integrin and CD151 on the surface of ECs using TIRFM (**Figure 6A-E**), followed by conditional colocalization analysis. For these experiments we used an antibody that recognizes total β1-integrin, regardless of its activation state (Byron et al., 2009) (in contrast to the positive control application above). Considering all pairs of target and reference objects (no conditional analysis yet), *p*(*TwR*) revealed significant overall colocalization between CD151 and β1-integrin (both ways), but not between CD36 and β1-integrin or CD36 and CD151 (**Figure 6F**). However, performing conditional colocalization analysis, and taking β1-integrin as the condition, the subset of CD36 colocalized with β1-integrin exhibited significant colocalization with CD151 (*p*(*TwR|TwC*) in **Figure 6G**). With CD151 as the condition, the subset of CD36 colocalized with CD151 exhibited slightly elevated, marginally significant, colocalization with β1-integrin (*p*(*TwR|TwC*) in **Figure 6H**; p-value = 0.1). Neither the fraction of β1-integrin nor the fraction of CD151 colocalized with CD36 was significant, regardless of the condition (**Suppl. Figure S3A, B**). This highlights the necessity for object-based colocalization analysis that allows for non-reciprocal colocalization relationships given the different abundance of different molecules. Finally, taking CD36 as the condition did not enhance the already significant colocalization between β1-integrin and CD151 (**Suppl. Figure S3C, D**).

**Figure 6:**
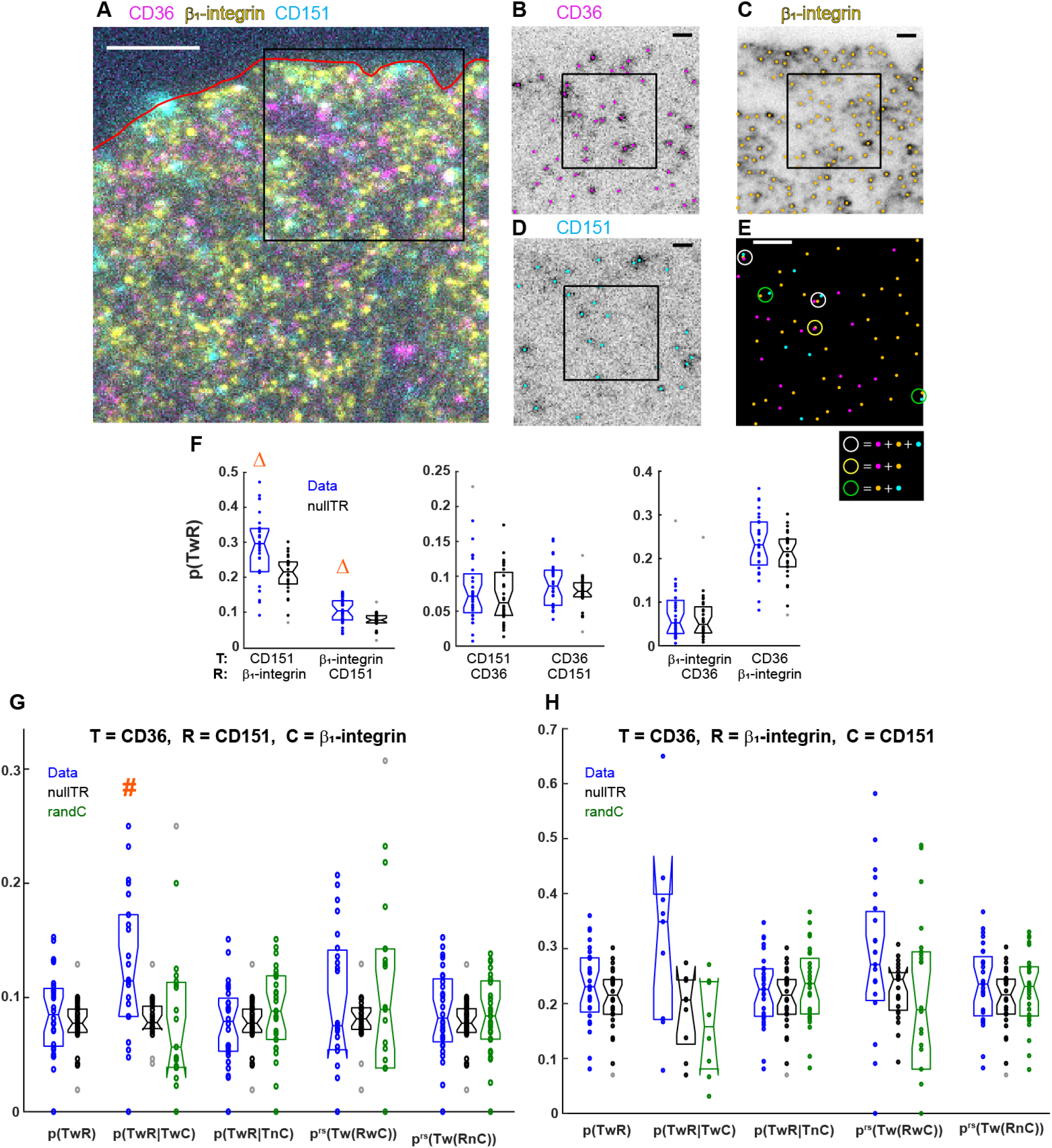
CD36 colocalizes with β1-integrin and CD151 together, with β1-integrin acting as the main mediator. **(A)** Representative three-color image of CD36, β1-integrin and CD151 on the surface of a TIME cell imaged via TIRFM. Red line shows cell edge. Scale bar, 5 μm. **(B-D)** Particle detections (shown as dots) overlaid on the individual channels for the area within the white box in (A). Scale bar, 1 μm. **(E)** Overlay of the three channel detections for the area within the black box in (B-D), with color-coding following that in (B-D). Colored circles (diameter = 267 nm) point out colocalization events between the different molecules, following color-coding in legend. Scale bar, 1 μm. **(F)** Overall probabilities of target to colocalize with reference (*p*(*TwR*)) for the different target and reference combinations among the imaged molecules, together with their coincidental counterparts (“nullTR”). Boxplots and dots as in Figure 3F. Orange triangles (Δ): p < 0.01, indicating that *p*(*TwR*) is significantly greater than its coincidental counterpart, as assessed via a Wilcoxon rank-sum test. **(G, H)** Conditional colocalization measures and their coincidental counterparts (“nullTR” and “randC”, as applicable) for indicated object combinations. Boxplots, dots and orange pound signs as in Figure 4G. *p^rs^*(*Tw*(*RwC*)|*TwC*) and *p^rs^*(*Tw*(*RnC*)|*TnC*) were omitted as in Figure 5G-J. N = 28 cells from 4 repeats. See **Table 3** for number of objects per channel and other dataset properties.

The CD36-independent colocalization between β1-integrin and CD151 is consistent with previous work that found direct association between β1-integrin and CD151 through their extracellular domains (Yauch et al., 2000). Our data suggest that CD36 interactions with β1-integrin bring CD36 to a complex that also contains CD151, through the latter’s interactions with β1-integrin. However, because the overall probability of CD36 colocalization with β1-integrin (*p*(*TwR*) in **Figure 6F**) is not significant, this suggests that the interaction of CD36 with β1-integrin also requires a mediator/stabilizer. This could be CD151, given the slight elevation of colocalization of CD36 with β1-integrin if CD36 is colocalized with CD151 (*p*(*TwR|TwC*) in **Figure 6H**), or some other molecule, or lipid nanodomain, not investigated in our study (Silverstein and Febbraio, 2009). The lack of significant pairwise colocalization between CD36 and β1-integrin could also be the result of labeling all β1-integrins in our study, regardless of the α-integrin partner, while CD36 may be interacting with particular αβ heterodimers (most likely α_6_β_1_ and/or α_3_β_1_ (Miao et al., 2001; Thorne et al., 2000)). Altogether, these analyses reveal that β1-integrin and CD151 colocalize together independently of CD36, while CD36 colocalizes with them together, as a complex, with β1-integrin acting as a major mediator/stabilizer between CD36 and CD151.

## Discussion

We have developed a new approach, conditional colocalization analysis, to assess the colocalization relationships and hierarchies between three molecular entities from multicolor microscopy images. Going beyond the question of whether two entities colocalize, it addresses the question whether the colocalization between two entities is increased or decreased by a third entity. We have demonstrated its specificity and sensitivity, and showcased its application to two different biological questions, one regarding the regulation of receptor-downstream effector interactions at specific plasma membrane locations, and one regarding the hierarchy of molecular interactions in multimolecular complexes. The source code for performing conditional colocalization analysis, along with detailed instructions for running it, is available at https://github.com/kjaqaman/conditionalColoc.

Conditional colocalization analysis of multicolor images offers an approach to shed light on interaction dependencies, hierarchies and regulation within the native, unperturbed environment of these interactions. It complements the classic approach of system perturbation to probe these questions in at least two ways. First, because it extracts information from the unperturbed system, it can provide unique insight in cases where molecular perturbations have side effects that complicate the interpretation of results, or where there is functional redundancy between molecules, in which case one molecule may compensate for another upon perturbation, thus masking the perturbed molecule’s true function(s) (Wagner, 2000; Zhang, 2012). Second, it enables the initial dissection of a system to gather evidence for the important players in regulating the interactions of interest, thus guiding later experiments involving molecular perturbations. Applying conditional colocalization analysis to super-resolution microscopy images instead of conventional microscopy images will bring the analyzed inter-object distances closer to molecular dimensions, thus deepening the insight that conditional colocalization analysis may provide about molecular interactions and their regulation (Huang et al., 2009; Sahl et al., 2017).

The information that conditional colocalization analysis of multicolor images yields about molecular interactions is difficult to obtain through other methods commonly employed to detect molecular interactions in their cellular context, such as Förster Resonance Energy Transfer (FRET) and Bimolecular Fluorescence Complementation (BiFC) (Romei and Boxer, 2019; Sun et al., 2013). Although some extensions have enabled the detection of triple interactions with FRET and BiFC (Kwaaitaal et al., 2010; Scott and Hoppe, 2015), these methods are mostly designed to yield signal only when the interaction of interest takes place. In other words, they primarily address the question of whether an interaction takes place or not. They do not allow the calculation of a probability of interacting, and thus they cannot be used to analyze the modulation of interactions without system perturbation.

While our application of conditional colocalization analysis has been focused on the molecular and macromolecular scale, this analysis approach is applicable to any biological system, as long as at least two out of the three imaged entities can be treated as punctate objects represented by their center coordinates. These objects do not have to be diffraction-limited; as long as they are approximately circular and of homogeneous size, their radius can be incorporated into the colocalization radius between objects. As for non-punctate objects, they can take any form or shape, as long as they are segmented properly, e.g. using ridge detection for curvilinear objects (Kittisopikul et al., 2020) or using deep learning approaches (Lucas et al., 2021). In addition, conditional colocalization analysis can be tailored to particular biological questions through question-specific selection of the objects analyzed (e.g. objects from a particular subcellular region, or objects of a certain size or shape). For example, our observation that β_1_-integrin at FAs does not show enhanced colocalization with pFAK hints at the transient residency of pFAK at FAs, which depends on the size of FAs (Kleinschmidt and Schlaepfer, 2017; Le Dévédec et al., 2012). Thus, it may be informative to analyze β_1_-integrin-pFAK colocalization in the context of different sizes of FAs.

Conceptually, conditional colocalization analysis can be extended in multiple ways. The same analysis, as currently implemented, can be applied to 2D live cell imaging, to determine the modulation of colocalization relationships over time, e.g. in response to a cellular stimulus. The analysis can be also readily extended to 3D, as long as there is sufficient axial resolution. Light-sheet microscopy with isotropic resolution would be ideally suited for this purpose (Power and Huisken, 2017). Finally, the analysis can be extended to analyze the colocalization relationships and hierarchies between more than three entities, as long as informative conditional probabilities and related measures can be defined. As extending the analysis to more than three colors will increase the total number of possible object combinations, the analysis procedure in this case may be iterative, to help identify biologically relevant object combinations. All in all, conditional colocalization analysis is a broadly-applicable method for the analysis of multicolor microscopy images. By dissecting the colocalization relationships between multiple molecular entities, it helps with the dissection of the complex landscape of multimolecular interactions in the cell.

## Materials and Methods

### Plasmids

A plasmid encoding CD36-fused at the N-terminus to HaloTag (Halo-CD36) was generated by PCR-cloning the amplification of HaloTag coding sequence (primers HaloTag-For and HaloTag-Rev detailed below (IDTDNA, Coralville, IA)) from pFN21K HaloTag® CMV Flexi® Vector (Promega, Madison, WI), followed by ligation in place of the mApple sequence in the mApple-CD36-C-10 vector (plasmid # 54874, Addgene, Watertown, MA (Githaka et al., 2016)), using restriction enzymes AgeI and BglII. Selected plasmid was verified by Sanger sequencing.

*HaloTag-For primer:* CATCCAGCGGGGATCCACCGGTCATGGCAGAAATCGGTACTGGCTTTCCATTCG
*HaloTag-Rev primer:* CCACTGCTGCCAGATCTCATGGCGATCGCGTTATCGCTCTG

### Cell culture and plating

Human Telomerase (hTERT)-immortalized microvascular endothelial cells isolated from human foreskin (TIME cells, ATCC, Manassas, VA) were grown in ATCC’s vascular cell basal medium supplemented with microvascular endothelial cell growth kit-VEGF and 12.5 μg/mL blasticidine (Sigma-Aldrich, St Louis, MO) for 48 h at 37°C + 5% CO_2_ until reaching 80-90% confluence. At confluence, cells were passaged and 5.4×10^4^ cells were plated on fibronectin-coated (10 μg/ml, MilliporeSigma, Burlington, MA), base/acid cleaned, 0.17 mm (#1.5) glass bottom dishes (14 mm glass diameter, MatTek, Ashland, MA) for 18 h prior to experiments.

### Transfection of Halo-CD36 in TIME cells

The Halo-CD36 plasmid was transiently transfected into TIME cells. Briefly, 5×10^6^ TIME cells (passages 14 to 21) were suspended in serum and antibiotic-free DMEM (ThermoFisher, Waltham, MA) and incubated with Halo-CD36 (1 μg) and UTR (Universal Transfection Reagent, Sigma) at 37°C for 20 min, following which the transfection mixture was added to a T-25 flask filled with complete DMEM medium. Transfected cells were plated on fibronectin-coated dishes, as described above, on day 1 post-transfection. Cells were imaged on day 2 post-transfection.

### Sample preparation for fluorescence imaging

For all samples except for CD36/β_1_-integrin/CD151, plated cells were washed once in wash buffer (HBSS + 1 mM HEPES and 0.1% NGS) to remove culture medium, and then incubated with HBSS containing or not 2 nM VEGF-A_165_ (Genscript, Piscataway, NJ) for 5 min at 37°C. Incubation solution was then removed and cells were fixed with a 3.2% Paraformaldehyde solution made in PBS (Electron Microscopy Sciences, Hatfield, PA) for 15 min at RT. Samples were then washed three times (5 min each) at RT with wash buffer, then permeabilized for 1 min with cold 0.01% Triton X-100 solution made in PBS. After three washes, samples were blocked for 15 min in blocking buffer (1% BSA, 5% NGS in wash buffer) and then incubated for 1 hr with primary antibodies (**Table 1**) at RT. After three washes, samples were incubated with secondary antibodies (**Table 1**) for 15 min at RT. After three more washes, they were then incubated with AlexaFluor647-conjugated primary antibodies (**Table 1**) for 15 min at RT. Finally, after three subsequent washes, dishes were incubated with imaging buffer (Oxyfluor 1%, Glucose 0.45%, Trolox 2 nM) in order to reduce photobleaching.

**Table 1.** Primary and secondary antibodies and their dilutions used for IF imaging.

| Antibody | Host, Clonality | Manufacturer (identifier) | Dilution used |
| --- | --- | --- | --- |
| Unconjugated primary |  |  |  |
| Anti- TfR | Rabbit, polyclonal | Abcam, ab84036 | 1:200 |
| Anti-TSAd (OTI4F3) | Mouse, monoclonal | Invitrogen, MA5-25894 | 1:100 |
| Anti-$\beta_1$-integrin (activated) (P4G11) | Mouse, monoclonal | Sigma-Aldrich, MAB1951 | 1:10000 |
| Anti-pFAK (pY397) (EP2160Y) | Rabbit, Monoclonal | Abcam, ab81298 | 1:10000 |
| Anti-VEGFR-2 (55B11) | Rabbit, Monoclonal | Cell Signaling Technology, #2479 | 1:400 |
| Anti-$\beta_1$-integrin (neutral) (K20) | Mouse, Monoclonal | Beckman Coulter, IM0791U | 1:75000 |

| Conjugated primary |  |  |  |
| --- | --- | --- | --- |
| Alexa Fluor® 647 Anti-CD151 (11G5a) | Mouse, Monoclonal | Abcam, ab199200 | 1:500 |
| Alexa Fluor® 647 Anti-Paxillin (Y113) | Rabbit, Monoclonal | Abcam, ab246719 | 1:100 |
| Alexa Fluor® 647 Anti-CHC (X22) | Mouse, Monoclonal | Invitrogen, MA1-065-A647 | 1:2000 |
| Secondary |  |  |  |
| Alexa Fluor® 488 Anti-Mouse IgG secondary | Goat, polyclonal | Invitrogen, A-11029 | 1:1000 |
| Alexa Fluor® 546 Anti-Rabbit IgG secondary | Goat, polyclonal | Invitrogen, A-11010 | 1:1000 |
| Alexa Fluor® 488 Anti-Rabbit IgG secondary | Goat, polyclonal | Invitrogen, A-11008 | 1:10000 |
| Alexa Fluor® 546 Anti-Mouse IgG secondary | Goat, polyclonal | Invitrogen, A-11003 | 1:10000 |

For CD36/β_1_-integrin/CD151, CD36 transfected cells (day 2 post-transfection) were rinsed with sterile DPBS and incubated with 50 nM JF549-Halo ligand (generous gift from Dr. Luke Lavis, Janelia Research Campus, Ashburn, VA) for 15 min in complete culture medium. This was followed by 3 quick washes in DPBS and then samples were incubated for 15 min in dye free complete culture medium. Incubations were at 37°C + 5% CO_2_. Following CD36 labelling, samples were washed once in wash buffer and fixed with a 4% Paraformaldehyde + 0.1% Glutaraldehyde (Electron Microscopy Sciences, Hatfield, PA) solution made in PBS, for 20 min at 4°C, in the dark. Freshly prepared quenching solution (0.1% NaBH4, Oakwood Chemical, Estill, South Carolina) was added immediately after fixation and cells were incubated for 7 min. Samples were then washed 3 times (5 min each) at RT with wash buffer. Samples were blocked for 30 min in blocking buffer and then incubated for 1 hr with a mouse primary antibody against β_1_-integrin at RT (**Table 1**). After three washes, samples were incubated with an AlexaFluor488-conjugated goat anti-mouse secondary antibody (**Table 1**) for 15 min at RT. After three more washes, they were then incubated with AlexaFluor647-conjugated primary antibody against CD151 (**Table 1**) for 30 min at RT. Finally, after three subsequent washes, dishes were incubated with imaging buffer before imaging.

### Total internal reflection fluorescence microscopy (TIRFM) imaging

For the TfR/TSAd/CHC, activated β1-integrin/pFAK/Pax, VEGFR2/TSAd/CHC and VEGFR2/TSAd/Pax datasets, cells were imaged at 37°C using an Olympus IX83 TIRF microscope equipped with a Z-Drift Compensator and a UAPO 100X/1.49 NA oil-immersion TIRF objective (Olympus, Center Valley, PA). The microscope was equipped with an iXon 888 1k × 1k EMCCD Camera (Andor - Oxford Instruments, Concord, MA). With an additional 1.6X magnification in place, the pixel size in the recorded image was 81 nm × 81 nm. Using the Olympus CellSens software, excitation light of 640 nm, 561 nm and 491 nm from an Olympus CellTIRF-4Line laser system was directed to the sample at a penetration depth of 90 nm by a TRF8001-OL3 Quad-band dichroic mirror. Fluorescence of different wavelengths was collected, filtered with emission filters of ET520/40m, ET605/52m and ET705/72m (Chroma, Bellows Falls, VT), and projected onto different sections of the camera chip by an OptoSplit III 3-channel image splitter (Cairn Research, Faversham, Kent, UK). The different channels were excited and recorded sequentially, in the order 640, then 561, then 491. Images were acquired with MetaMorph (Molecular Devices, San Jose, CA). Camera EM gain was set to 100 for all acquisitions.

For all molecule combinations mentioned above except for activated β_1_-integrin/pFAK/Pax, each channel was imaged once using an exposure time of 99 ms and a laser power of 3.6, 6.4 and 5.9 mW at the coverslip (measured with a Thorlabs power meter at an incident angle of 0°) for 491 nm, 561 nm and 640 nm, respectively. As for the activated β_1_-integrin/pFAK/Pax combination, even with the high antibody dilutions employed, the β_1_-integrin and pFAK images were not always punctate. To get reliably punctate images, β_1_-integrin and pFAK were imaged in streaming mode for 300 frames (at 10 Hz), with laser power increased to 5.4 mW for the 491 channel and 9.6 mW for the 561 channel, in order to photobleach the signal. The last frame of each stream, where the β_1_-integrin and pFAK signals were largely punctate, was then used as the image for β_1_-integrin and pFAK detection and colocalization analysis. Paxillin (the 640 channel) was imaged only once, as with the other molecule combinations.

For the CD36/β_1_-integrin/CD151 dataset, cells were imaged at 37°C using an S-TIRF system (Spectral Applied Research, via BioVision Technologies, Exton, PA) mounted on a Nikon Ti-E inverted microscope with Perfect Focus and a CFI Apo 60X/1.49 NA oil-immersion TIRF objective (Nikon Instruments, Melville, NY), equipped with an Evolve EMCCD camera (Photometrics, Tucson, AZ). A custom 3X tube lens was employed to achieve an 89 nm × 89 nm pixel size in the recorded image. Sequential illumination by a 488 nm diode laser (Coherent), a 561 nm diode pumped solid state laser (Cobolt), and a 637 nm diode laser (Coherent) was achieved through an ILE laser merge module (Spectral Applied Research), with 488 nm, 561 nm and 637 nm laser powers of 3.65, 13.2 and 10.6 mW, respectively, at the coverslip (measured with a Thorlabs power meter at an incident angle of 0°). The penetration depth was set to 100 nm via the Diskovery platform control (Spectral Applied Research). Three-channel images were acquired with MetaMorph (Molecular Devices, San Jose, CA), with 100 ms exposure per image, using an EM gain of 100. Emission filters of ET520/40m, ET605/52m and ET705/72m (Chroma, Bellows Falls, VT) were employed.

For every three-channel image or movie, a brightfield snapshot of the imaged cell region was also acquired, in order to aid with manual delineation of the region of interest (ROI) mask for the ensuing analysis.

#### Computing environment

All image and data analysis and simulation tasks were performed in Matlab R2020a (The MathWorks, Natick, MA). All employed code and software packages are compatible with Matlab R2020a on a Linux 64-bit operating system. Images were loaded into Matlab using Bio-Formats (Linkert et al., 2010).

### Punctate object detection

Punctate objects were detected using the “point-source detection” particle detection algorithm in u-track (https://github.com/DanuserLab/u-track) (Aguet et al., 2013; Jaqaman et al., 2008). In brief, the algorithm consists of two steps: (i) a filtering step to determine pixels likely to contain objects, and (ii) a Gaussian fitting step to determine the object positions with sub-pixel localization. With the appropriate, wavelength-dependent standard deviation, a two-dimensional Gaussian is a good approximation of the microscope’s point spread function (Thomann et al., 2002; Zhang et al., 2007). Largely default parameter values were used, except for the alpha-value for determining object detection significance by comparing the fitted Gaussian amplitude to the local background noise distribution (**Table 2**). The alpha-value was chosen based on visual assessment of the detection results, with the goal of minimizing both false positives (superfluous detections) and false negatives (missed particles). The number of objects detected within the ROI for each dataset is listed in **Table 3**.

**Table 2.** Three-channel imaging and detection of indicated molecule combinations. PSF, point spread function. Shown are the parameters of “point-source detection” that vary between channels and datasets. Two other parameters that change (namely “Max Fit Adjust” and “Fit Window Size”) are not shown because they were given their default values of 2×PSF sigma and 4×PSF sigma, respectively. The paxillin (Pax) channel has N/A parameters because it was not detected using point-source detection due to its non-punctate nature.

|  | Channel wavelength | PSF sigma (pixels, nm) | Parameters |
| --- | --- | --- | --- |
| **TfR/TSAd/CHC** |  |  |  |
| TfR | 561 | 1.4, 113 | Alpha: 0.01, all others: default. |
| TSAd | 488 | 1.2, 97 | Alpha: 0.01, all others: default. |
| CHC | 640 | 1.58, 128 | Alpha: 0.05, all others: default. |
| **activated β1-integrin/pFAK/Pax** |  |  |  |
| activated β1-integrin | 561 | 1.4, 113 | Alpha: 0.01, all others: default. |
| pFAK | 488 | 1.2, 97 | Alpha: 0.01, all others: default. |
| Pax | 640 | N/A | N/A |
| **VEGFR-2/TSAd/CHC** |  |  |  |
| VEGFR-2 | 561 | 1.4, 113 | Alpha: 0.05, all others: default. |
| TSAd | 488 | 1.2, 97 | Alpha: 0.01, all others: default. |
| CHC | 640 | 1.58, 128 | Alpha: 0.05, all others: default. |
| **VEGFR-2/TSAd/Pax** |  |  |  |
| VEGFR-2 | 561 | 1.4, 113 | Alpha: 0.05, all others: default. |
| TSAd | 488 | 1.2, 97 | Alpha: 0.01, all others: default. |
| Pax | 640 | N/A | N/A |
| **CD36/β1-integrin/CD151** |  |  |  |
| CD36 | 561 | 1.4, 113 | Alpha: 0.01, all others: default. |
| β1-integrin | 488 | 1.2, 97 | Alpha: 0.01, all others: default. |
| CD151 | 640 | 1.58, 128 | Alpha: 0.05, all others: default. |

**Table 3.** Number of objects per ROI, ROI area and segmented patch area (where applicable) for the various experimental datasets.

| Dataset<br>(Channels 1/2/3 or<br>561/488/640) | Channel 1<br>objects per<br>ROI<br>[min max]<br>avg | Channel 2<br>objects per<br>ROI<br>[min max]<br>avg | Channel 3<br>objects per<br>ROI<br>[min max]<br>avg | ROI area (pixels)<br>[min max]<br>avg | Segmented<br>patch area<br>(pixels)<br>[min max]<br>avg |
| --- | --- | --- | --- | --- | --- |
| TfR/TSAd/CHC | [151 660]<br>368 | [80 506]<br>214 | [233 751]<br>490 | [85,190 219,960]<br>156,800 | N/A |
| $\beta_1$-Integrin/pFAK/<br>Pax | [157 804]<br>371 | [424 1,561]<br>910 | [47 178]<br>101 | [61,570 234,760]<br>132,950 | [87 159]<br>114 |
| VEGFR-2/TSAd/ Pax | [357 1,194]<br>754 | [174 663]<br>358 | [8 218]<br>135 | [93,880 231,610]<br>158,340 | [73 140]<br>96 |
| VEGFR-2/TSAd/<br>CHC | [352 1,134]<br>695 | [200 639]<br>415 | [284 928]<br>568 | [62,510 202,080]<br>144,870 | N/A |
| CD36/$\beta_1$-Integrin/<br>CD151 | [34 914]<br>192 | [191 1,871]<br>715 | [61 739]<br>254 | [27,760 209,170]<br>92,783 | N/A |

### Non-punctate object segmentation

Paxillin patches (“non-punctate objects”) were segmented using an intensity threshold to separate foreground from background. The threshold was determined for each image individually. It was taken as the 90^th^ percentile of each image’s intensity distribution after noise filtering (using a Gaussian kernel with standard deviation = 1 pixel) and local background subtraction. Local background was estimated for each image by filtering it with a Gaussian kernel with standard deviation = 10 pixels. The threshold was then applied to the noise-filtered and background-subtracted image to segment the paxillin patches. Patches with an area of at least 30 pixels were retained for further analysis. The number of segmented patches within the ROI and their areas are listed in **Table 3**.

### Region of interest (ROI) mask segmentation

ROI masks, covering the imaged area of each cell of interest, were delineated manually. For TfR/TSAd/CHC, VEGFR-2/TSAd/CHC and CD36/β1-integrin/CD151, delineation was based on the brightfield image. For VEGFR-2/TSAd/Pax and beta1 integrin/pFAK/Pax, delineation was based on the brightfield image as well as paxillin and pFAK (where imaged).

### Core algorithm to calculate colocalization fraction of target with reference

Our implementation closely followed the original algorithm proposed in (Helmuth et al., 2010). In brief, for punctate target and reference objects, the distance between any two objects was defined as the Euclidean distance between their estimated positions (centers). For punctate target and non-punctate reference objects, the distance was defined as the Euclidean distance between the estimated position (center) of the target object and the closest pixel in the segmentation of the non-punctate reference object. With this, a target object was considered to be colocalized with a reference object if the distance between it and its nearest neighbor reference object was smaller than a user-defined threshold (“colocalization radius”). The colocalization radius accounted for various experimental considerations, such as registration shifts between channels, segmentation accuracy, etc. The fraction of target colocalized with reference (*f_coloc_*) was then calculated as the ratio of the number of target objects colocalized with reference objects to the total number of target objects.

To calculate the coincidental colocalization fraction of target with reference 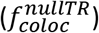 given the cellular context, in particular the number and spatial distribution of reference objects within the ROI mask, we followed (Helmuth et al., 2010) and replaced the target positions with a grid of positions covering the whole ROI. Specifically, we used each pixel within the ROI as one grid point (i.e. target position), and repeated the above distance calculations, classification and fraction calculation.

### Position randomization of condition objects within the ROI mask

To calculate the conditional colocalization probabilities expected by chance, i.e. in the absence of any true influence of the condition objects on target-reference colocalization, the condition objects were randomized within the ROI mask. In the case of punctate condition objects, to ensure that none of the randomly chosen locations were too close to the ROI mask boundary, the ROI mask was first eroded using a square structuring element with size equal to the colocalization radius plus three pixels (i.e. six pixels in our applications). The new condition object locations were then chosen within the eroded ROI mask, using the same number of condition objects as in the original.

In the case of non-punctate condition objects, we conserved not only the number of objects, but also the shape of each object. To optimize the selection of random locations, condition objects in this case were placed in random locations sequentially, starting with the largest object, and going down to the smallest object. The idea behind this order was that it was easier to place small objects between large objects than the other way around. Once an object was placed, the pixels it occupied were removed from the ROI mask, as those pixels were no longer available for any other object (**Suppl. Figure S1**). In addition, before placing each object, the mask was eroded using a disk structing element with radius equal to the average radius of the object being placed (**Suppl. Figure S1**). The purpose of this was to avoid placing any new object too close to already-placed objects (leading to overlap) or to the ROI mask boundary (leading to part of the object being outside the ROI mask). With this strategy, we were able to randomize the locations of non-punctate condition objects within the ROI mask while avoiding any overlap between them, and while avoiding the scenario of not enough contiguous space to place larger objects (a scenario that happened often when smaller objects were placed before larger objects).

### Validation simulations

As mentioned in Results, the validation simulations were performed within cell ROI masks taken from our experimental data, to keep the simulations as realistic as possible. Each mask was used as a repeat for each parameter combination. Cell masks were eroded using a square element with side length = 10 pixels to avoid molecules being positioned too close to the cell edge.

#### Punctate target objects and non-punctate reference objects

To simulate non-punctate reference objects, the paxillin patch segmentations belonging to the employed cell masks were used as reference objects. Then, based on the input *p*(*TwR*), target objects were divided into colocalized or not colocalized with reference objects. Target objects colocalized with reference objects were placed 0-2 pixels from the boundaries of the reference objects with which they were associated. Target objects not colocalized with reference objects were placed at random coordinates within the eroded cell mask, excluding the pixels belonging to reference objects. For this, the reference objects were dilated using a square element with side length = 10 pixels to exclude the colocalization area around each reference object when placing not-colocalized target objects.

#### Punctate target and reference objects

These simulations followed the strategy described above for non-punctate reference objects. The only difference was that they did not use the experimental paxillin patch segmentations for reference objects. Instead, reference objects were placed at random coordinates within the eroded cell mask, and a binary image was created where pixels of value one represented the positions of reference objects. These masks were the starting point for the simulations as described for the previous case.

#### Punctate target and reference objects, with non-punctate condition objects

As with the case above of non-punctate reference objects, the paxillin patch segmentations belonging to the employed cell masks were used for the simulations, but this time as condition objects. Based on the input *p*(*TwC*) and *p*(*RwC*), target and reference objects were divided into condition-positive and condition-negative groups. Condition-positive reference objects were placed 0.2 pixels away from the boundaries of the condition objects with which they were associated. This very close placement was motivated by practical considerations; it made it easier to place target objects (as described next) close to both a reference object and a condition object when needed, as being placed close to one almost guaranteed being placed close to the other. Condition-negative reference objects were placed at random coordinates within the eroded cell mask, excluding the pixels belonging to condition objects after dilating them using a square element with side length = 10 pixels.

To place target objects, they were subdivided further. First, based on *p*(*TwR*|*TwC*), the condition-positive target objects were divided into colocalized or not colocalized with reference objects. Condition-positive target objects colocalized with reference objects were placed 0-2 pixels from condition-positive reference objects. Given the placement scheme for condition-positive reference objects, this almost guaranteed that the target objects were colocalized with condition objects as well. Condition-positive target objects not colocalized with reference objects were placed 0-2 pixels from the boundaries of condition objects, after eliminating the boundary pixels associated with reference objects. Second, based on *p*(*TwR*|*TnC*), condition-negative target objects were divided into colocalized or not with reference objects. Condition-negative target objects colocalized with reference objects were placed 0-2 pixels from condition-negative reference objects, which largely placed them far from condition objects as well. Condition-negative target objects not colocalized with reference objects were placed at random coordinates within the eroded cell mask, excluding the pixels belonging to condition objects and reference objects, both after dilation using a square element with side length = 10 pixels.

#### Punctate target, reference and condition objects

These simulations followed the strategy described above for non-punctate condition objects. The only difference was that they did not use the experimental paxillin patch segmentations for condition objects. Instead, condition objects were placed at random coordinates within the eroded cell mask, and a binary image was created where pixels of value one represented the positions of condition objects. These masks were the starting point for the simulations as described for the previous case.

### Derivation of expected colocalization measures from simulation input parameters

In our simulations, *p*(*TwC*), *p*(*RwC*), *p*(*TwR*|*TwC*) and *p*(*TwR*|*TnC*) were explicitly defined as input parameters. For validation, their values as calculated by our conditional colocalization analysis were compared directly to their input values. The remaining two conditional colocalization measures, *p^rs^*(*Tw*(*RwC*)) and *p^rs^*(*Tw*(*RnC*)), and the overall colocalization probability *p*(*TwR),* were not explicitly defined. Thus, for validation, their calculated values were compared to their expected values, as derived from the input *p*(*TwC*), *p*(*RwC*), *p*(*TwR*|*TwC*) and *p*(*TwR*|*TnC*).

The expected value of *p*(*TwR)* was calculated from the input parameters in a straightforward manner using the law of total probability:

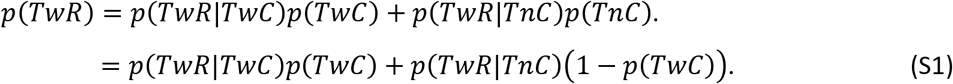

To calculate the expected values of *p^rs^*(*Tw*(*RwC*)) and *p^rs^*(*Tw*(*RnC*)), we assumed one simplifying relationship between objects (**Assumption 1**), an assumption that was satisfied in our simulations (see above).

##### Assumption 1

If a target object and a reference object are colocalized, then either both are colocalized with a condition object, or neither is colocalized with a condition object.

With **Assumption 1**, and given the relationships between the different target subgroups (**Figure 1C, D**), it followed that:

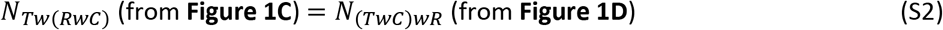

Using the definitions of *p*(*Tw*(*RwC*)), *p*(*TwR*|*TwC*) and *p*(*TwC*) (Eqs. 1, 6 and 8), it then followed that:

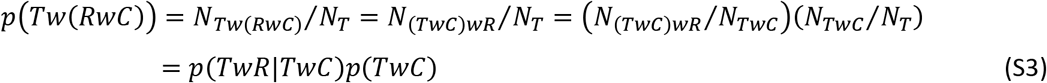

Using Eq. S3 in Eq. 10, the expected value of *p^rs^*(*Tw*(*RwC*)) could thus be calculated from the input parameters as follows:

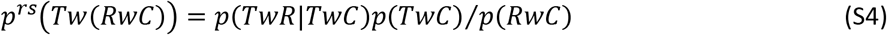

With similar arguments, the expected value of *p^rs^*(*Tw*(*RnC*)) was calculated from the input parameters as follows:

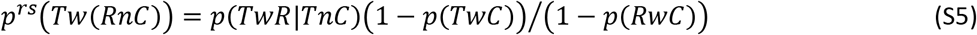

## Supplemental Material

There are three supplemental figures.

## Acknowledgements

We thank Dr. Tieqiao Zhang for microscopy support, Dr. Anthony R. Vega for initial programming of code for pairwise colocalization of punctate objects, and the Lavis lab (Janelia Research Campus, Ashburn, VA) for JF549-Halo ligand. This work was supported by funding from NIH/NIGMS (R35 GM119619), the Welch Foundation (I-1901) and the UT Southwestern Endowed Scholars Program to KJ, and the Canadian Institutes of Health Research (CIHR PS 165816) and Natural Sciences and Engineering Research Council (NSERC RGPIN-2018-05783) to NT. JVL was an honorary trainee of the NIH Molecular Biophysics Training Grant (5T32GM131963; PI: Dr. Yuh Min Chook), and was also supported by a Diversity Supplement to NIH grant R35 GM119619 (PI: KJ). The authors declare no competing financial interests.

## Author contributions

KJ, JVL and BRA designed research; JVL and KJ developed algorithm; JVL wrote software; BRA and AD performed experiments; JVL and BRA performed analysis; NT made HaloTag-CD36 plasmid; KJ, JVL and BRA wrote paper with input from all co-authors.

---

**Suppl. Figure S1:**
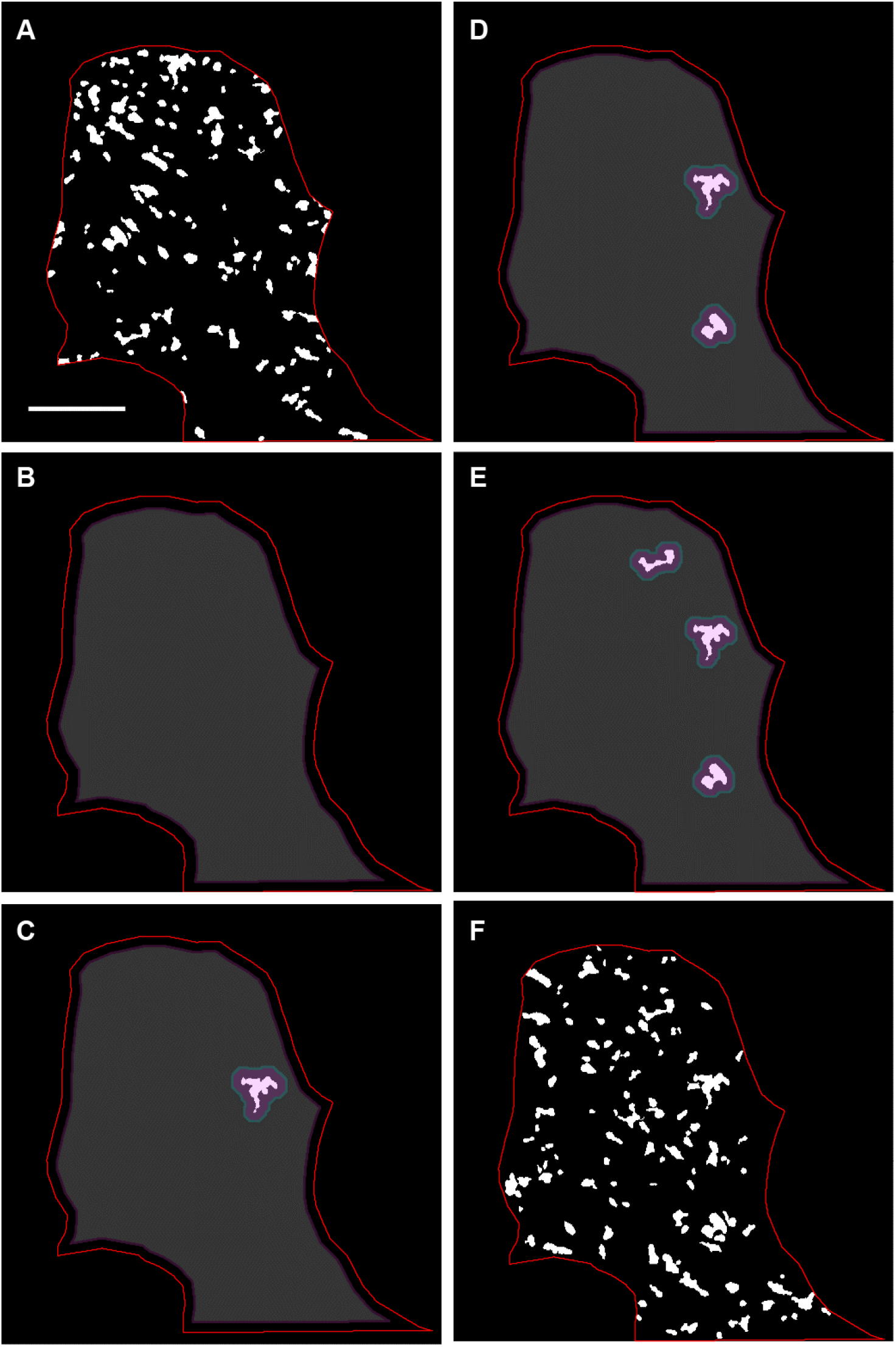
Example illustrating randomization procedure for non-punctate condition objects. **(A)** Binary image of original paxillin patch segmentation. Red line outlines region of interest (ROI; the cell in the image in this case). Scale bar (applicable to all panels), 10 μm. **(B)** Available area for randomly placing the first condition object (gray-shaded area) within total ROI. **(C)** Random placement of first (largest) condition object, and the remaining available area (gray-shaded area), which excludes an area around the already placed object. **(D, E)** Random placement of second largest (D) and third largest (E) condition objects, and the remaining available area after each placement (gray-shaded area). **(F)** Final random placement of condition objects within the ROI.

**Suppl. Figure S2:**
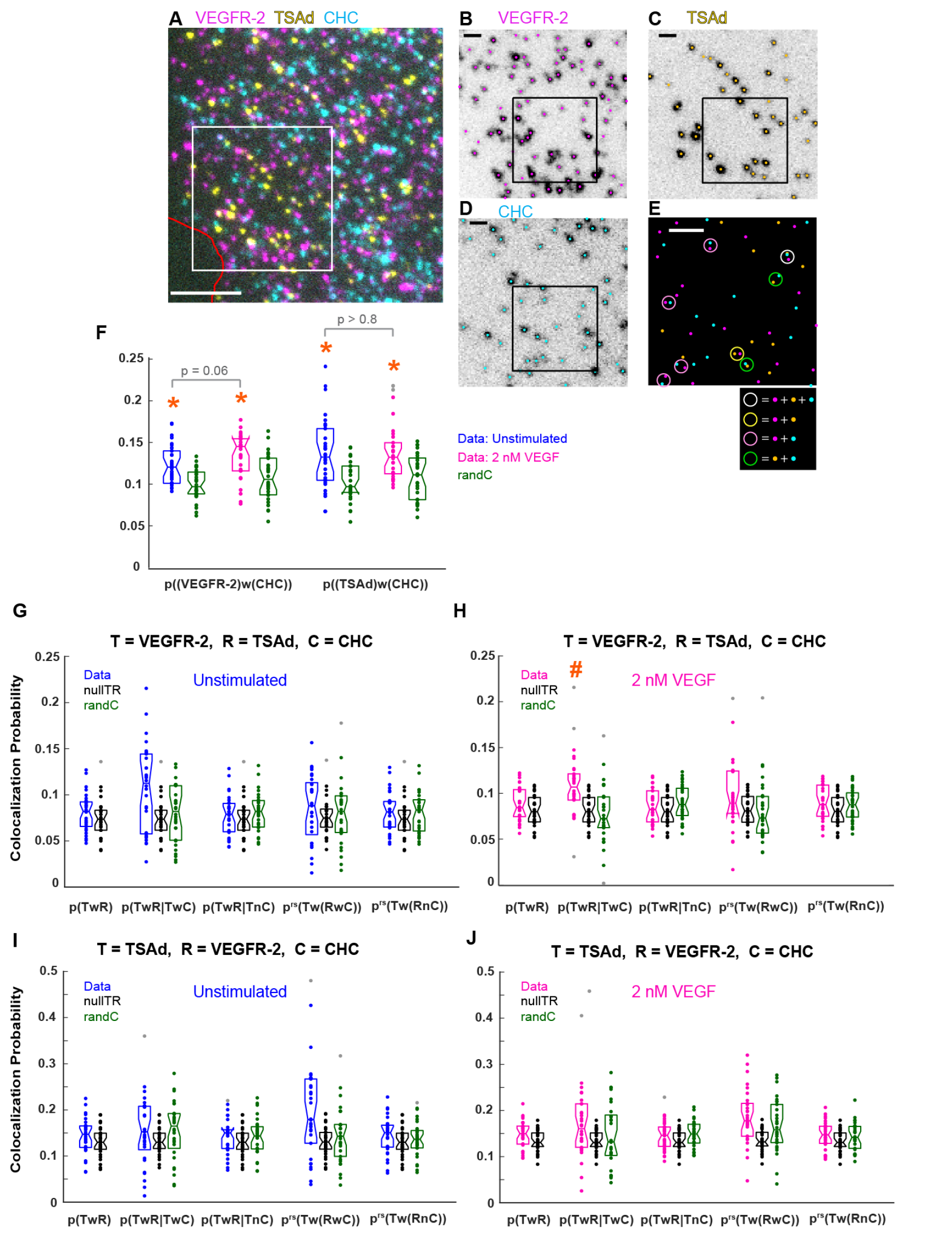
VEGFR-2-TSAd colocalization is minimally influenced by their localization at CCSs. **(A)** Representative three-color IF image of VEGFR-2, TSAd and CHC on the surface of a TIME cell imaged via TIRFM. Red line shows cell edge. Scale bar, 5 μm. **(B-D)** Particle detections (shown as dots) overlaid on the individual channels for the area within the black box in (A). Scale bar, 1 μm. **(E)** Overlay of the three channel detections for the area within the black box in (B-D), with color coding following that in (B-D). Colored circles (diameter = 243 nm) point out the colocalization events between the different molecules, following color-coding in legend. Scale bar, 1 μm. **(F)** Probabilities of VEGFR-2 and TSAd to colocalize with the condition (CHC; representing CCSs), in the presence or absence of VEGF, together with their coincidental counterparts (“randC”). Boxplots, dots and orange asterisks as in Figure 3F. The probabilities in the absence and presence of VEGF were compared to each other using a Wilcoxon rank-sum test, resulting in indicated p-values. **(G-J)** Conditional colocalization measures and their coincidental counterparts (“nullTR” and “randC”) for target = VEGFR-2 and reference = TSAd **(G, H)** and vice versa **(I, J)**, both with condition = CHC, in the absence **(G, I)** or presence **(H, J)** of 2 nM VEGF. Boxplots, dots and orange pound signs as in Figure 4G. *p^rs^*(*Tw*(*RwC*)|*TwC*) and *p^rs^*(*Tw*(*RnC*)|*TnC*) were omitted as in Figure 5G-H. As no measure was significant in both the absence and presence of VEGF, no statistical tests were performed to compare the two. N = 30 cells from 3 repeats. See **Table 3** for number of objects per channel and other dataset properties.

**Suppl. Figure S3:**
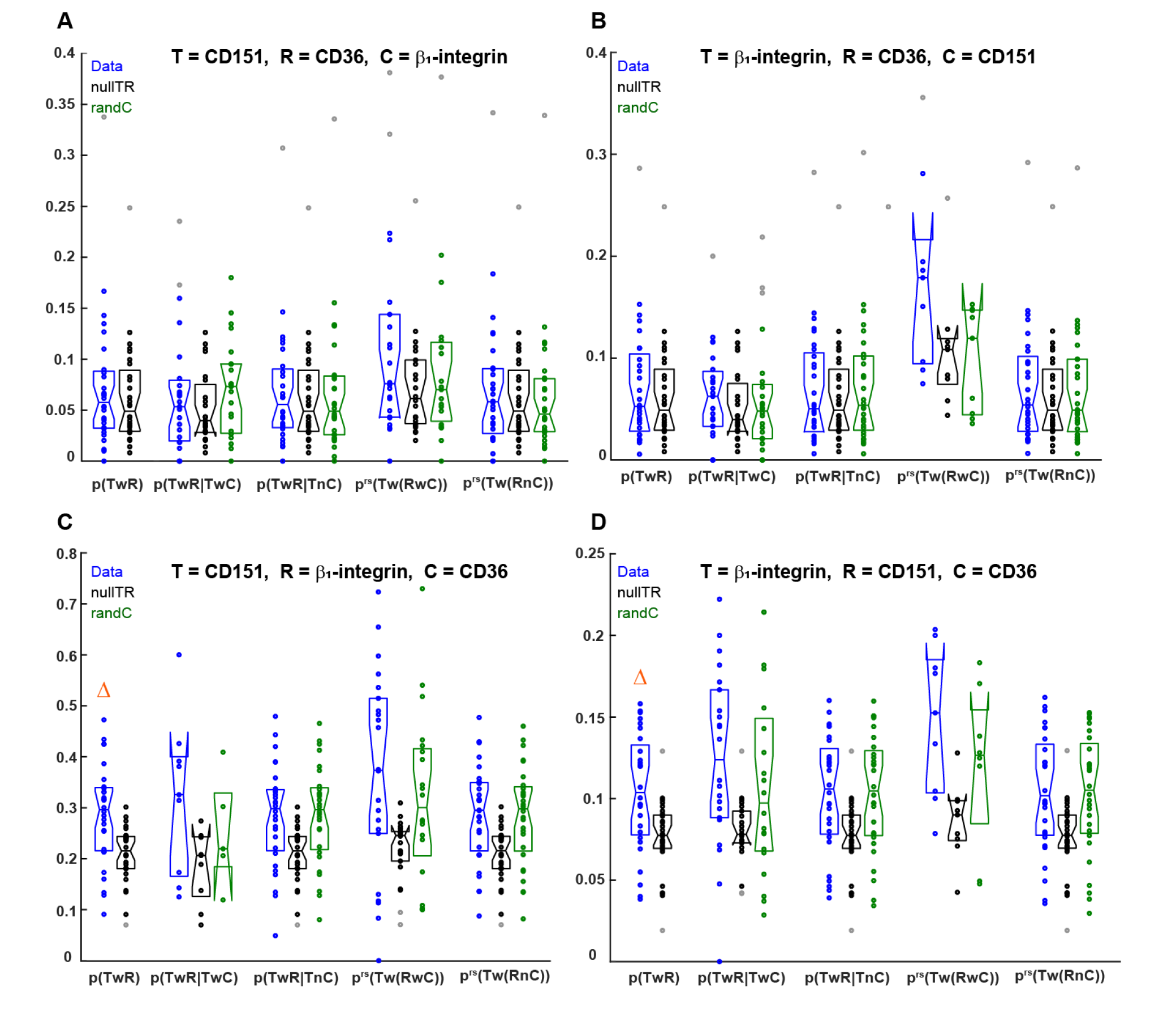
Conditional colocalization analysis of CD36, β1-integrin and CD151 for the target, reference and condition permutations not shown in Figure 6. **(A-D)** Conditional colocalization measures and their coincidental counterparts (“nullTR” and “randC”) for the following target, reference and condition combinations: **(A)** Target = CD151, reference = CD36 and condition = β1-integrin (target and reference switched relative to Figure 6G). **(B)** Target = β1-integrin, reference = CD36 and condition = CD151 (target and reference switched relative to Figure 6H). **(C)** Target = CD151, reference = β1-integrin and condition = CD36. (**D)** Target = β1-integrin, reference = CD151 and condition = CD36 (target and reference switched relative to (C)). Boxplots, dots and orange triangles as in Figure 6F. Other than *p*(*TwR)* in (C) and (D), none of the conditional colocalization measures pass the significance test, and thus none are highlighted. N as in Figure 6. See **Table 3** for number of objects per channel and other dataset properties.

